# Molecular pathology of acute spinal cord injury in middle-aged mice

**DOI:** 10.1101/2025.05.08.652873

**Authors:** Corey Fehlberg, Danny John, Brian Kang, James S. Choi, Susana Cerqueira, Alexis Brake, Jae K. Lee

## Abstract

The median age of sustaining a spinal cord injury has steadily increased from 29 to 43 over the last several decades. Although more pre-clinical studies in aged rodents are being done to address this shift in demographics, there have not been comprehensive transcriptomic studies investigating SCI pathobiology in middle-aged mice. To address this gap in knowledge, we compared behavioral, histopathological, and transcriptional outcomes in young (2-4 months-old) and middle-aged (10-12 months-old) mice. In contrast to previous studies, open field tests showed no differences in locomotor recovery between the young and middle-aged mice over a one-month period. The injury site also demonstrated similar histopathology in terms of lesion size, and numbers of macrophages and fibroblasts. Acutely after injury, proliferation of macrophages, fibroblasts, and astrocytes were also similar between the two age groups. In addition, spatial transcriptomics showed similar transcriptionally defined regions around the injury site at 3 days post injury. However, single cell RNA-sequencing of the cells at the injury site and surrounding spared tissue showed differences in select cell subpopulations. Taken together, our results indicate that although young and middle-aged mice display similar locomotor recovery and histopathology after SCI, changes in cell subpopulations may underly a decline in repair mechanisms that are manifested after this age.

## Background

Contusive spinal cord injury (SCI) ranks with the most devastating forms of trauma by morbidity, often resulting in long-term disability due to the limited regenerative capacity of the central nervous system (CNS). Numerous factors complicate the clinical management of SCI, with advanced age at the time of injury being notable among the least favorable prognosticators [1, 2]. Importantly, the life-point at which SCI is sustained is known to play a crucial role in clinical recovery, as patients under 20 exhibit better outcomes than counterparts 65 years and older [3, 4]. Similar findings have been reported in young (2-4 month-old) and aged (18-24 month-old) murine SCI studies [5–7]. The median age of incidence for SCI has shifted from young (twenties) to middle-aged (forties), which according to the Jackson Labs, is equivalent to 10-14 months old in mice. Although a limited number of previous studies have also reported worse functional and pathological outcomes in middle-aged mice [7–10], molecular pathology using comprehensive transcriptomic approaches have not been performed. Thus, our goal in this study was to use spatial and single cell transcriptomics to dissect the pathobiology of SCI in middle-aged mice.

In addition to injury site pathology, axon regeneration promoted by genetic deletion of PTEN is robust in young mice but diminishes with age [11]. However, regenerative capacities of CNS axons might be tract-specific, and axons such as serotonergic axons maintain their regenerative capacity even in old age [12]. Similar age-related decline in regeneration has been reported after peripheral nerve injury [13, 14]. While some of this loss in regenerative capacity is due to neuron-intrinsic mechanisms, the injury environment is also often more inflammatory in older animals [11, 15]. Previous studies on middle-aged mice and rats have shown worse behavioral recovery and injury site pathology in mice and rats [7–10]. The Gensel lab has shown that middle-aged mice show higher reactive oxygen species at the injury site, which makes them better responders to antioxidant treatment after SCI compared to young mice [8]. Middle-aged rats have also shown worse functional recovery as well as greater demyelination after SCI [7]. Collectively, these findings underscore the reality of age-related declines in SCI recovery as demonstrated in multiple rodent models.

Contusive SCI in rodents follow a fairly stereotypic acute pathobiology. On the first day post-injury (1 dpi), the inflammatory response by neural cells such as microglia and astrocytes amplifies the innate immune response through chemotactic recruitment of peripheral myeloid cells (primarily neutrophils and monocytes) to the injury site. By 3 dpi, most glial cells, including astrocytes, microglia, and oligodendrocyte progenitor cells (OPCs), peak in their proliferative activity, initiating gliosis. Meanwhile, monocytes differentiate into various macrophage phenotypes to both engulf cellular debris and recruit perivascular fibroblasts which, in turn, trigger fibrosis. This cell proliferation gives rise to a diverse cellular subpopulation that provide valuable insight into cellular heterogeneity and lineage after SCI [16]. These and other pivotal factors make this timepoint an especially target-rich environment for investigating SCI pathobiology.

In this study, we investigated the potential differences in SCI pathobiology between young and middle-aged mice. We employed a combination of behavioral assessments, histological analyses, and single-cell RNA sequencing (scRNA-seq) and spatial transcriptomics, allowing for a comprehensive investigation of processes across multiple levels of biological organization ranging from tissue architecture of the injury site to gene expression patterns in the individual cell. Contrary to previous studies, we did not find significant differences in gross histopathology, or locomotor recovery between young and middle-aged mice. Spatial transcriptomics also did not reveal major differences in spatial organization of the injury site between the two age groups. However, scRNA-seq revealed changes in specific myeloid, glial, and vascular cell subpopulations in middle-aged mice, which may underly the behavioral deficits previously reported in aged rodents and humans after SCI.

## Methods

### Spinal cord injury surgeries and locomotor assessment

Contusive spinal cord injury (SCI) was performed as previously described [17]. Female C57BL/6J (Jackson Labs, stock 000664) mice aged 2-3 months (young) or 10-12 months (middle-aged) were anesthetized with a combination of ketamine (100 mg/kg) and xylazine (15 mg/kg) administered intraperitoneally (number of animals are noted in each experiment below). Following anesthesia, T8 laminectomy was performed to expose the mid-thoracic spinal cord, and the spinal column stabilized with spinal clamps on the adjacent T7 and T9 vertebrae. A contusion injury was then induced using an Infinite Horizon Impactor device (Precision Systems and Instrumentation, LLC), delivering a force of between 65 and 70 kilodynes. Post-operative care included the subcutaneous administration of slow-release buprenorphine (3.25 mg/kg) for analgesia injected immediately after surgery. Lactated Ringer’s solution (1 ml) and gentamicin (5 mg/kg) were injected subcutaneously once immediately after surgery and once daily for three days following surgery to prevent dehydration and infection. Bladders were manually expressed twice daily until the conclusion of the experiment. Locomotor recovery was assessed by two people using the Basso Mouse Scale open field test for mice [18]. Animals were scored at day 1 post-SCI and weekly thereafter (n=10 Young, n=9 middle-aged). Scores for left and right hindlimbs were averaged for each animal at each time point. Scorer were blinded to the experimental groups. All procedures adhered to the guidelines of the University of Miami Institutional Animal Care and Use Committee and the National Institutes of Health.

### Tissue Processing for Spatial Transcriptomics

On the day of tissue collection, animals were deeply anesthetized with avertin (250 mg/kg, i.p) and then transcardially perfused with ice-cold PBS (n=2 per group). A 4mm section of the spinal cord centered at the injury site was dissected, blotted dry with kimwipe, and embedded in OCT for parasagittal cryosectioning. Tissue blocks were stored at −80 °C until sectioning. Tissue permeabilization time was determined in accordance with the Visium Tissue Optimization (TO) user guide (10x Genomics CG000238 Rev E). Briefly, 10 µm thick tissue sections were placed on the TO slide. One capture area was left empty as a negative control and 2 ug bulk-extracted spinal cord RNA (Qiagen RNeasy plus kit) was added to another empty well as a positive control. Following cDNA synthesis and tissue removal, fluorescent cDNA footprints were visualized using Nikon Eclipse microscope and 12 minutes was identified as the optimal permeabilization time.

Prior to sectioning, the tissue blocks and Visium Spatial Gene Expression slide were equilibrated to the cryostat chamber for 30 minutes. To correctly identify the injury epicenter, we sampled the tissue after every eighth 10 µm section on a separate slide. Myelin was stained for 5 minutes with 1% Sudan Black in propylene glycol pre-warmed to 56°C in a water bath and washed with running tap water until the water ran clear [19]. A 10 µm tissue section toward the midline of the tissue and containing a large injury site was captured on the Spatial Gene Expression slide. These steps were repeated for a total of four biological replicates (one per capture area) on a single slide. The full slide was transported on dry ice to a thermocycler fitted with the Visium thermocycler adapter (3000380, 10x Genomics) pre-warmed to 37°C. After thawing on the thermocycler adapter for 1 minute, the slide was fixed in ice-cold methanol at −20°C for 30 minutes. Myelin was again stained with 1% Sudan Black in propylene glycol for 5 minutes at room temperature. The excess staining solution was discarded, and the slide was carefully immersed 3 times in a 50 mL conical tube of ultrapure water, then 15 times each in two additional beakers with 800 mL ultrapure water. The slide was cover slipped with 85% glycerol mounting media according to the manufacturer’s instruction and imaged using Nikon Eclipse inverted microscope using brightfield microscopy (Supp. Fig. 2A).

After imaging, the Spatial Gene Expression slide was immersed in 3X SSC at a 45°angle to allow the coverslip to gently separate from the slide. The tissue was enzymatically permeabilized for 12 minutes as determined by the above tissue optimization experiment and cDNA synthesis and library preparation was performed following the manufacturer’s guidelines (Visium Spatial Gene Expression Reagents Kits User Guide CG000160, Rev F). cDNA amplification cycle number was determined by the Cq (defined by 25% peak RFU) value obtained by qPCR.

### RNA-sequencing for spatial transcriptomics

Each spatial sequencing sample consisted of a single animal per 10X Visium V1 sequencing area. The cDNA library generated was sequenced on an Illumina NovaSeq with a SP-200 flow cell for paired-end sequencing targeting 50 million reads per sample. Sequences were aligned and filtered with the mm10-2020-A genome using Space Ranger V2.0.0. Passing reads recorded as over tissue were then loaded into R and processed as a Seurat object.

### Analysis of spatial transcriptomics data

Counts were normalized with SCTransform, regressing by sample, and clustered using the standard Seurat pipeline, without spatial information [SCTransform (assay = ‘Spatial’, vars.to.regress = ‘SampleName’, variable.features.rv.th = 1.3), FindNeighbors (reduction = ‘pca’, dims = 1:10), FindClusters (resolution = 10), RunUMAP (reduction = ‘pca’, dims = 1:10)]. Sequences and spots were not filtered by percent mitochondrial or ribosomal counts, or by number of UMIs detected. The resulting clusters were then annotated by their differentially expressed genes, Gene Ontology terms, and anatomical locations after mapping the spots back to the original tissues. Gene expression differences between young and aged tissues for each layer were found by pseudo bulking by sample and layer, then running FindMarkers with default settings for each layer between ages. Gene ontology information was obtained by passing the top differentially expressed genes for each layer to the Ensembl database with current ‘org.Mm.eg.db’ mappings, using the fisher test for significance, then excluding any terms with less than 5 genes mapped.

### Tissue Processing for Single Cell RNA-seq

Mice used for single cell sequencing were processed as previously described [16]. Briefly, each mouse was anesthetized with Avertin (250 mg/kg i.p.) prior to transcardial perfusion with a pre-oxygenated artificial cerebrospinal fluid (aCSF) solution. From each, an 8-mm section of the spinal cord centered at the injury site was then dissected and placed in ice-cold cutting solution upon extraction until the final dissection. For every 8 mm spinal cord (approximately 400 mg), 50μL of Enzyme P dissolved in 1.9 mL of buffer X was prepared. The solution was then pre-warmed for at least 10min in 37°C water bath. After peeling away the meninges, the parenchymal tissue was finely chopped 10-15x in horizontal and vertical directions with a razor blade, washed with 5 ml of cutting solution, and centrifuged at 300 g for 2 minutes at RT. The resulting cell pellet was processed using the Miltenyi Neural Tissue Dissociation Kit-P (cat #130–092–628) following the manufacturer’s protocol. The cell suspensions were manually triturated (slowly, 10 times) using a large-opening (1,000 μm diameter) fire-polished pipette. After a 10-minute incubation at 37°C, the suspensions were further triturated with medium- and then small-opening (750 μm and 500 μm diameter, respectively) fire-polished pipettes to create a single-cell suspension. The suspensions were then filtered through a 70 μm cell strainer into a 50 mL conical, washed with 10 ml of HBSS with Ca^2+^/Mg^2+^, and centrifuged at 4°C and 300 g for 5 minutes. The supernatant was aspirated, and the cell pellets were resuspended in 1mL ACK Lysis Buffer for 1 minute at RT, immediately followed by the addition of 5 mL HBSS with Ca^2+^/Mg^2+^, and then centrifuged at 300 g for 5 minutes at 4°C, and the supernatant was discarded. Samples were then processed using Miltenyi Myelin Removal Beads according to manufacturer guidelines. The injury site from a separate mouse was processed as above followed by Miltenyi anti-ACSA-2 magnetic beads to enrich for astrocytes (keeping the cells retained in the column and discarding the flowthrough). The resulting cell suspension was then centrifuged at 300 g for 5 minutes at 4°C, and the supernatant was discarded. One non-enriched and one astrocyte-enriched samples were pooled to generate one biological replicate.

### RNA-sequencing for single cell transcriptomics

Each single cell sequencing sample consisted of two pooled animals with one sample enriched for astrocytes. The eight biological replicates consisted of one young uninjured, one young 3 days post-injury (dpi), three middle-aged uninjured, and three middle-aged 3dpi samples. Prior to sequencing, cell suspensions were assessed for purity, concentration, and viability with AOPI Stain imaged on a Nexcelom K2 Cellometer. After confirming viability was above 80%, cells were diluted when necessary to achieve approximately 10,000 cells per-capture. ∼7,000 cells were sequenced per-sample with a mean of ∼95k reads per-cell and a mean of ∼11k UMIs per-cell. Sample libraries were prepared with the 10X Chromium Next GEM Single Cell 3’ Reagent Kits v3.1 (Dual Index) and multiplexed on the sequencer with Dual Index Kit TT (Set A) according to manufacturer’s protocols. These libraries were then sequenced using paired-end sequencing on eight channels of an Illumina Novaseq X 10-billion-read flow cell with a 300-cycle kit. The resulting sequences were then aligned and filtered using the CellRanger 6.0.2 pipeline. Briefly, reads from FASTQ files were aligned to the mm10-3.0.0 reference genome. Using the STAR splice-aware aligner, the reads were aligned and excluded if they mapped to <50% of an exonic sequence or contained single base substitution errors, then UMIs were removed. The final count matrix was generated containing all passing UMIs in each droplet.

### Analysis of single cell RNA-seq data

Droplets were initially processed using DropletUtils’s emptyDrops function and barcode-ranking to identify droplets which likely did not contain cells. The droplets were further processed to remove probable low-quality cells using sample-level metrics. Droplets with scores outside of three median absolute deviations of the median were considered low quality for the log values of the total UMI counts and number of unique features. Cells were further removed if their mitochondrial count percentage exceeded 30%; our lab has found that lower thresholds disproportionately exclude monocytes. Putative cell quality metrics before and after filtering are present in supplemental figure 1A and 1B, respectively. To manually exclude nondescript cells and include monocytes, crude clustering using Seurat’s recommended pipeline [NormalizeData, FindVariableFeatures, ScaleData, RunPCA (npcs = 20), FindNeighbors (dims = 1:20), FindClusters (resolution = 1.3), RunUMAP (dims = 1:20)] was performed on each sample and clusters were manually annotated for filtering. After droplets were processed as above, all passing cells in each sample were then processed using Scrublet. Doublet rates were set using estimated rates published by 10X. Flagged cells were confirmed through clustering and comparing known marker genes. Downstream in the analysis if segregating clusters showed overlap in known marker genes between the three meta groups (i.e. *Aqp1* and *Rgs1*) they were removed. Cells with overlap in known marker genes within the three meta groups (i.e *Pecam* and *Rgs1*) were not excluded. After processing each sample individually, samples were combined and only genes represented in all datasets were retained.

Raw data matrices of counts by droplets generated for each sample were then individually processed as follows. Matrices were filtered for empty drops using a combination of barcode UMI ranking to remove droplets which fall below the inflection point and the emptyDrops function from the DropletUtils package [20, 21] (setting the lower bounds to the lower knee of the UMI ranking and retaining points above the inflection point). The resulting cells were then filtered on generalized quality control metrics [16] (<30% mitochondrial genes, <3 MAD in both log10 counts and log10 features). Calculated doublets were then labeled with Scrublet [22] using estimated doublet rates provided by 10X. Predicted cells were then manually checked by coarse clustering of each sample using Seurat’s default pipeline (with nfeatures = 2000, npcs = 20, dims = 1:20, resolution = 0 to 6). Resulting clusters were then checked for expression of known marker genes [16] and any expressing elevated markers from cells of two different niches (myeloid, vascular, glial) were removed; cells expressing markers for two cell types of the same nice were retained (i.e macrophages and neutrophils or endothelial cells and pericytes). The resulting cells were then re-clustered and checked until sufficiently cleaned (Supp. Fig. 1C). Any doublet groups which appeared in downstream processing were similarly checked and removed. Resulting samples were then combined without regression by metadata with our previously published dataset [16] clustered (with nfeatures = 2000, npcs = 20, dims = 1:20, resolution = 2), and annotated for broad cell types. This results in a total of 4 young uninjured, 3 young 3dpi, 3 middle aged uninjured, and 3 middle aged 3dpi samples. Samples retrieved from GSE162610 are labeled as Milich and contain all young samples. Young and middle-aged samples generated in this study are labeled as such. We used a combination of expression of known marker genes [16], filtering using scGate [23], and mapping to existing datasets with SingleR [24]. The annotated cellular clusters were then split by niche, re-clustered, and re-annotated by cell subtype as before. Identities were confirmed using marker genes and by gene ontology of top differentially expressed genes.

Differential gene expression tests were performed on matrices at the single-cell and pseudobulk levels. Distinct homeostatic and injury-responsive subtypes were compared. Cells without distinct pre- and post-injury subtypes were compared between uninjured and 3dpi in the same group (BAMs, B-cells, T-cells, Fibroblast-1, Pericytes, and VSMCs). Ependymal and Endothelial cell subtypes were pooled for their respective homeostatic states. We used the FindMarkers and FindAllFeatures Seurat functions with default settings (unless otherwise stated) to perform the Presto Wilcoxon Rank Sum test with Bonferroni correction on cells grouped by comparison. Pseudobulk gene matrices were generated using the AggregateExpression Seurat function, grouping by sample name and cell type. Genes retrieved from FindMarkers were filtered to an adjusted p-value of < 0.001, and all other genes in the global environment were set to 1. This list was then analyzed using the topGO package for biological processes (statistic = ‘fisher’, algorithm = ‘classic’) and filtered for the top 20 terms.

### Immunohistochemistry and quantification

Samples were processed as previously described [25]. Mice were anesthetized with avertin and transcardially perfused with ice-cold 4% paraformaldehyde (PFA) in PBS. Whole spinal cords were dissected, post-fixed in 4% PFA for 2 hours, then transferred to a 30% sucrose solution in PBS for cryoprotection (1-3 days). An 8-mm segment of thoracic spinal cord centered on T8 was isolated, embedded in O.C.T. compound (Tissue-Tek), and cryo-sectioned at a thickness of 16 μm. The serial tissue sections were mounted across 10 Superfrost Plus slides and stored at −20°C. On the day of immunohistochemistry, sections were thawed and dried at room temperature (RT) and washed with PBS three times for 5 min each at RT. Sections were then incubated with 5% Normal Donkey Serum in PBST (0.3% TritonX-100 in PBS) for 45 min at RT. The sections were then incubated with 5% Normal Donkey Serum in PBST with primary antibodies overnight at 4°C (see below for antibodies and concentrations). The next day, sections were washed with PBST three times for 5 min each at RT. Sections were incubated with secondary antibodies in PBST for 60 min, DAPI (1:50,000) for 5 min, then PBST three times for 5 min each all at RT. Glass coverslips were mounted using Fluoromount-G (Southern Biotech; cat #0100-01) and sealed using CoverGrip (Biotium; cat #23005). Primary antibodies used: rat anti-CD11b (1:200, BioRad, MCA74GA), chicken anti-GFAP (1:500, ab4674, Abcam), rabbit anti-PDGFRβ (1:200, ab32570, Abcam). Secondary antibodies used were Donkey anti-Rabbit Alexa Fluor 488 (1:500), Donkey anti-Chicken Alexa Fluor 568 (1:500), Donkey anti-Rat Alexa Fluor 647 (1:500). Quantification of tissue sections was performed using ImageJ-Fiji software (version 2.9.0). For all analyses, images were acquired using an Olympus VS120 Slide Scanner with a 20X objective. At least three sections centered on the injury site were quantified and averaged per-animal (one biological replicate). Contours were drawn outlining the GFAP^-^ region for lesion area. The perilesional area was defined as 250 μm surrounding the GFAP^-^ area. A grid was placed over the injury site and every third box fully within the contour was quantified for cell proportions. 28 dpi CD11b (macrophages) and PDGFRβ (fibroblasts) density was quantified by counting the number of CD11b^+^/DAPI^+^ or PDGFRβ^+^/DAPI^+^ cells, respectively, out of all DAPI^+^ nuclei in the lesion and normalized to the lesion area. 4 dpi CD11b and PDGFRβ proliferation was quantified by counting the number of CD11b^+^/EDU^+^/DAPI+ or PDGFRβ^+^/EDU^+^/DAPI^+^ cells, respectively, out of all CD11b^+^/ DAPI^+^ or PDGFRβ^+^/DAPI^+^ cells in the lesion and normalized to the lesion area. The same procedure was repeated for the perilesional space and normalized to the perilesional area. Astrocyte proliferation (GFAP^+^/EDU^+^/DAPI^+^ cells out of all GFAP^+^/DAPI^+^ cells) was only quantified in the perilesional space.

### Statistical Analysis

Quantitative data from tissue analysis were analyzed using GraphPad Prism (version 9.5.0). Statistical comparisons between groups were made using unpaired Student’s t-tests or one-way ANOVA, followed by post-hoc tests where appropriate. Cell proportions were compared globally at the subtype level with a two-way ANOVA with a Tukey test for multiple comparisons. A p-value of <0.05 was considered statistically significant.

## Results

### Young and middle-aged mice display similar locomotor recovery and histopathology after SCI

We first evaluated locomotor recovery using the BMS scale at 1, 7, 14, 21, and 28 dpi. Both young and middle-aged cohorts exhibited pronounced hind-limb paralysis acutely after injury, with incremental recovery over time. Importantly, there were no statistical differences between the two age groups at any timepoint on either the main BMS score (Fig. 1A) or the subscore (Fig. 1B) during the first month post-injury. We also compared the histopathology at 28 dpi between the two age groups by immunostaining for GFAP (astrocytes), CD11b (macrophages-myeloid cells), and PDGFRβ (fibroblasts). Our results show that the lesion size (as determined by the GFAP-negative area) was similar between the two age groups (Fig. 1C-E). In addition, the density of macrophages (Fig. 1F-H) as well as fibroblasts (Fig. 1I-K) were also similar between young and middle-aged injury sites. Taken together, our data indicate that locomotor recovery and histopathology between young and middle-aged mice are similar chronically after contusive SCI.

**Figure 1.**
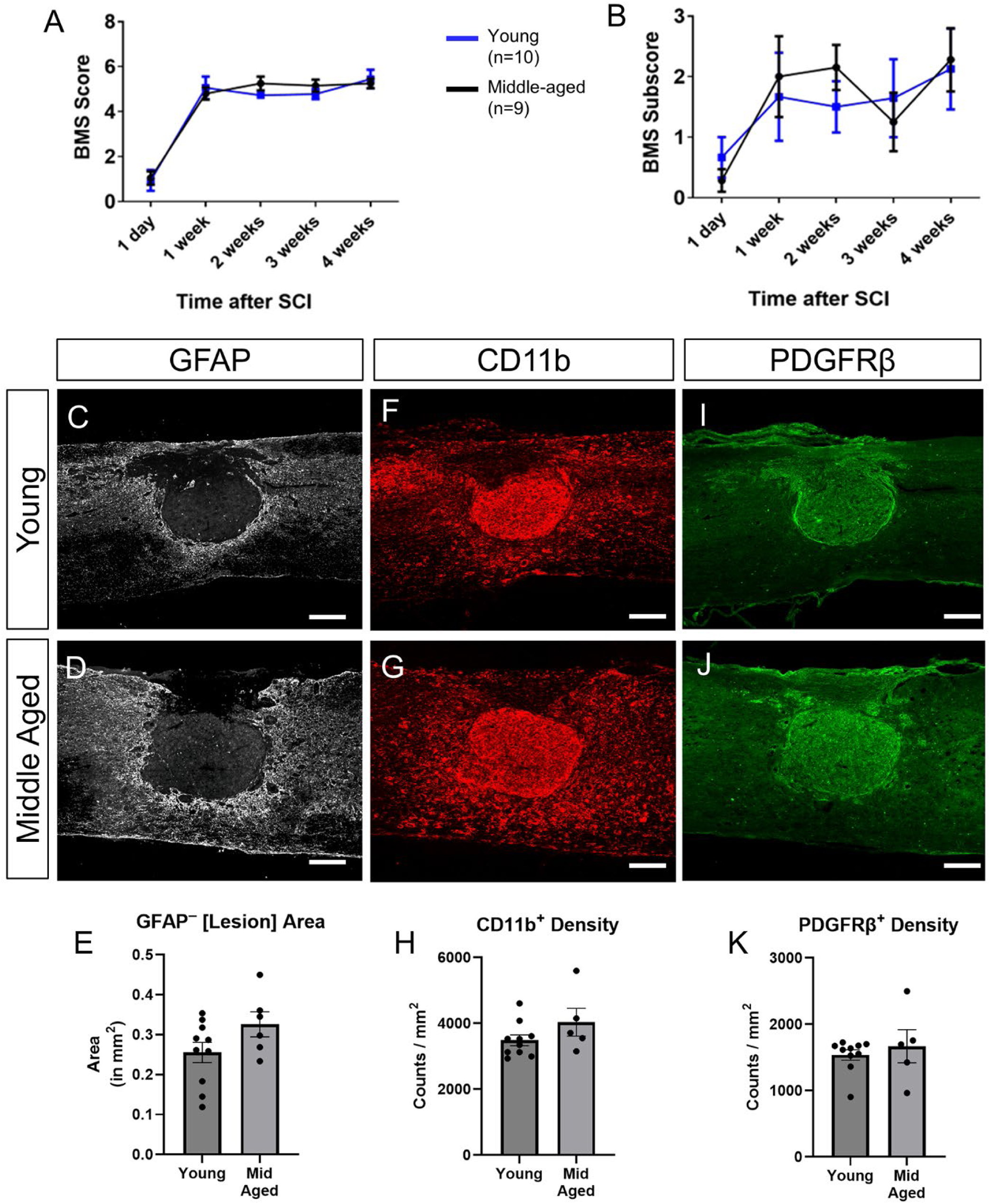
Young and middle-aged mice show similar locomotor recovery and lesion pathology at 4 weeks after SCI. (A) Basso Mouse Scale (BMS) scores and (B) subscores of young (n=10) and middle-aged (n=9) mice over a 4-week period post-spinal cord injury (SCI). Two-way repeated measures ANOVA with Bonferroni post-test. n=9 for young and n=10 for middle aged mice. Representative images of GFAP+ (gray C,D), CD11b (red F, G), and PDGFRβ (green, I, J) immunostaining in spinal cord lesion areas of young and middle-aged mice. (E) Quantification of the GFAP-negative lesion area did not show differences between the two groups. (H) Quantification of the density of CD11b+ myeloid cells in the lesion area did not show differences between the two groups. (K) Quantification of the density of PDGFRβ+ fibroblasts in the lesion area did not show differences between the two groups. Data are presented as mean ± SEM. Unpaired t-test, each data point is a biological replicate. Scale bar = 200 µm.

To assess whether cell proliferation at the injury site is affected in middle-aged mice, we injected EdU at 3 dpi to label proliferating cells and histologically assessed the injury site at 4 dpi. By co-immunostaining for EdU and GFAP, CD11b, or PDGFRβ, we quantified the number of proliferating astrocytes (Fig. 2A-C), macrophages (Fig. 2D-F), and fibroblasts (Fig. 2G-I) respectively. Our data did not show any differences between young and middle-aged mice in the proliferation of any of the three cell types examined. In addition, similar to our findings in the 28 dpi tissue above, the lesion size as well as the total number of macrophages and fibroblasts were not different between the two age groups even at this acute time point (Fig. 2J-L). Taken together, our data indicate that neither the proliferation of nor the number of astrocytes, macrophages or fibroblasts are different between young and middle-aged SCI mice.

**Figure 2.**
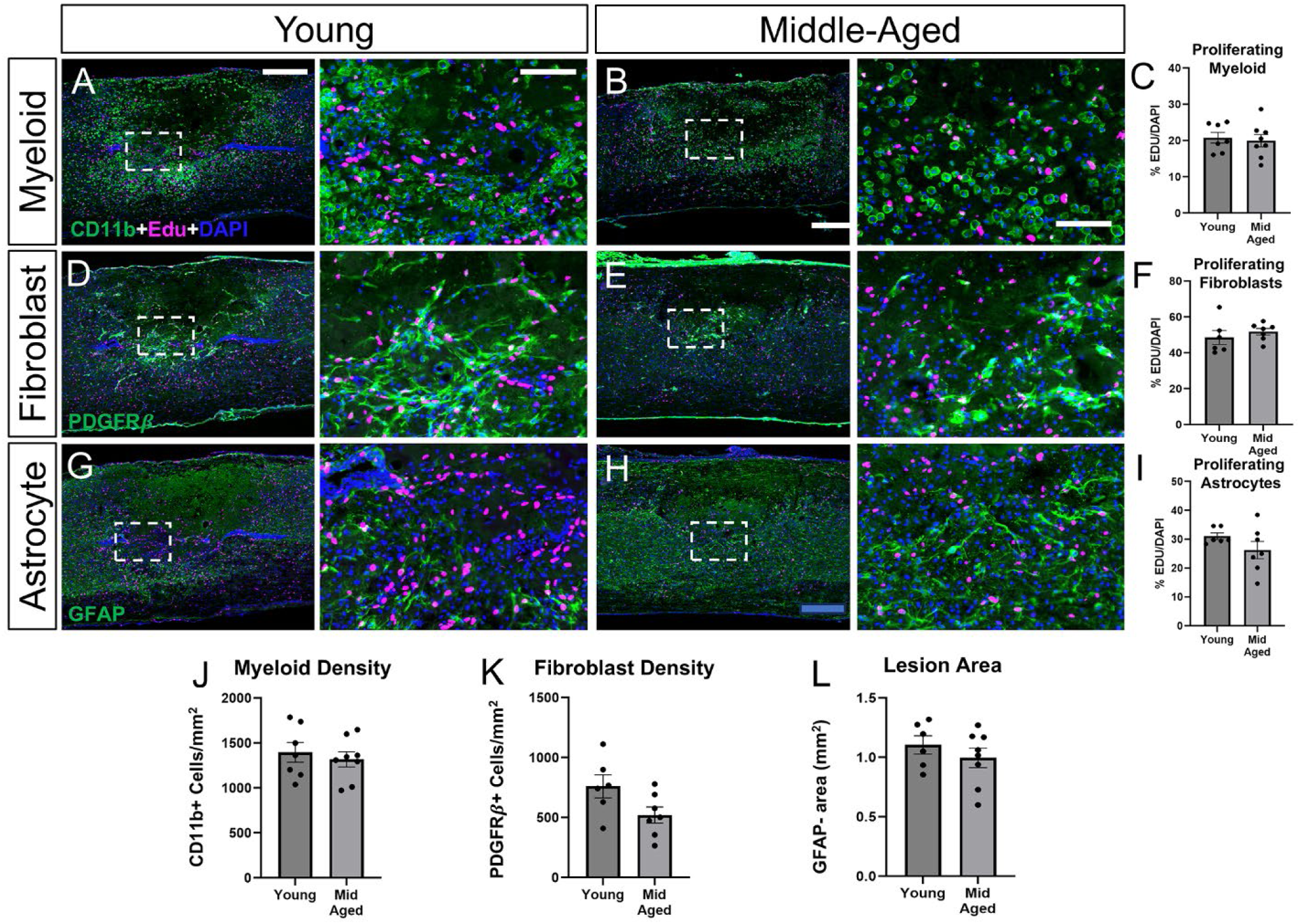
Young and middle-aged mice show similar cell proliferation and lesion pathology acutely after SCI. (A, B) Representative images of CD11b+ myeloid cells (green) co-stained with EdU (magenta) in young (A) and middle-aged (B) mice at 4dpi. (C) Quantification of CD11b+/EdU+ cells as a percentage of all CD11b+ cells within the GFAP-negative lesion area. (D, E) Representative images of PDGFRβ+ fibroblasts (green) co-stained with EdU (magenta) in young (D) and middle-aged (E) mice at 4 dpi. (F) Quantification of PDGFRβ+/EdU+ cells as a percentage of all PDGFRβ+ cells within the GFAP-negative lesion area. (G, H) Representative images of GFAP+ astrocytes (green) co-stained with Edu (magenta) in young (G) and middled-aged (H) mice at 4 dpi. (I) Quantification of GFAP+/Edu+ cells as a percentage of all GFAP+ cells within the perilesional region (250 μM region bordering the lesion). (J, K) Density of CD11b+ myeloid cells (J) or PDGFRβ+ fibroblasts (K) within the lesion area in young and middle-aged mice. (L) Area of the GFAP-negative lesion in young and middle-aged mice. All counted cells were DAPI+ (nucleus in blue). Error bars = SEM. Each data point is a biological replicate. Unpaired Student’s t-test. Boxed region is enlarged in the adjacent image on the right. Scale bar = 400µm for low magnification images, and 100µm for high magnification images.

### Spatial transcriptomics shows similar injury site domains between young and middle-aged mice acutely after SCI

To gain a much more comprehensive insight into the molecular histopathology of young and middle-aged mice after SCI, we performed spatial transcriptomics using the 10X Visium platform on two young and two middle-aged 3 dpi mice. Clustering of the barcoded spots pertaining to each tissue section resulted in 8 domains that defined the injury site and the surrounding spared tissue. The injury site domains were defined as the infiltrating injury core, necrotic, scar interface, and stromal layer. The surrounding spared tissue domains were defined as white matter, dorsal horn, gray matter, and ventral motor neurons (Fig. 3A, Supp. Fig. 2B). To identify these regions, we used a combination of annotated genes, gene ontology terms based on top differentially expressed genes, as well as their spatial location (Fig. 3B, C). Notably, the injury-associated layers were more similar to each other in their broad transcriptional profiles (Pearson Correlation Coefficient, 2000 genes, Δr>0.5) than compared to the spared tissue domain, likely due to their non-neural nature.

**Figure 3.**
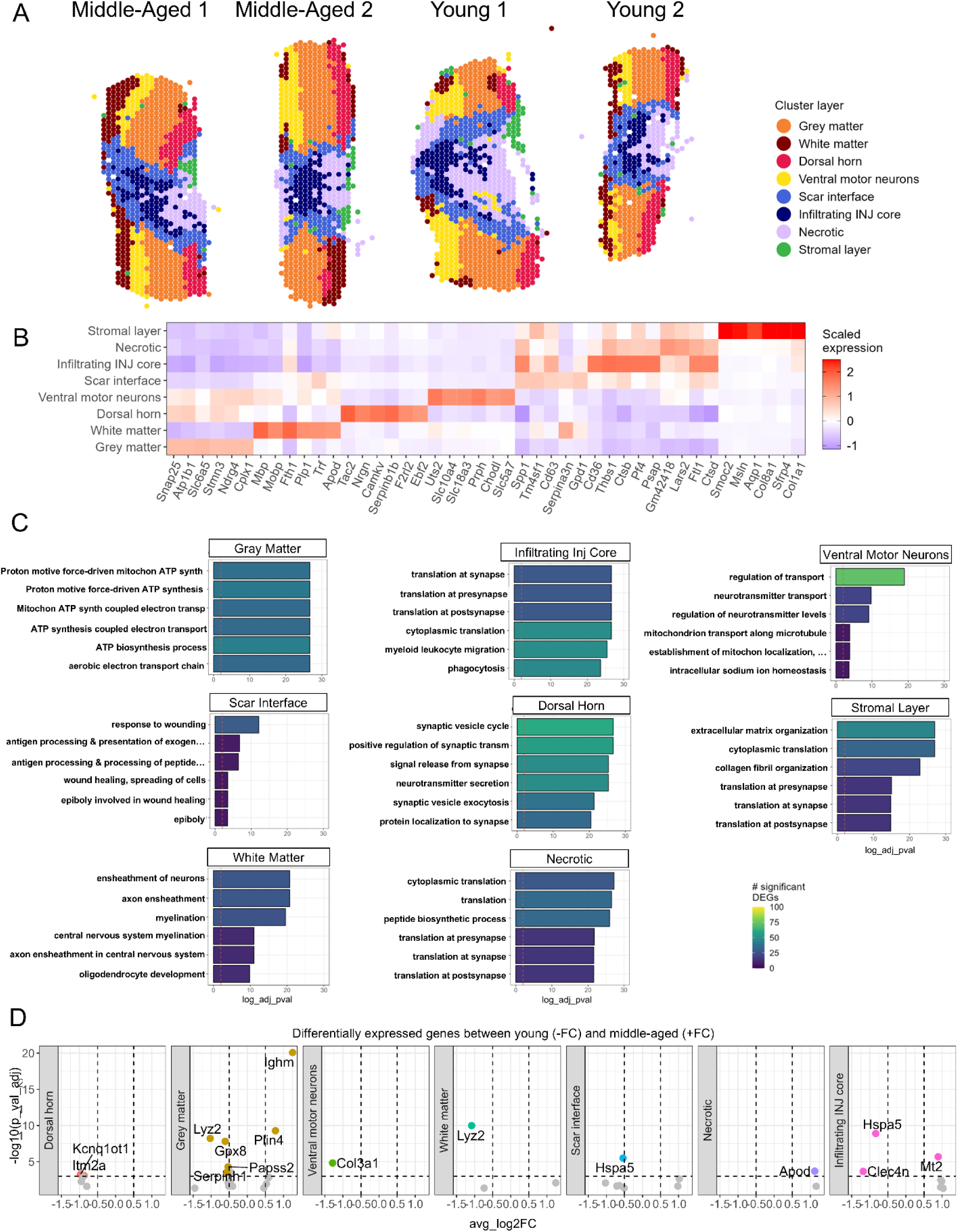
Spatial transcriptomics identifies distinct domains in and around the injury site that are similar between young and middle-aged mice. (A) 10X Visium spatial transcriptomic spots were clustered based on gene expression, and mapped on the injury site to identify eight distinct spatial domains that were named based on a combination of marker genes, Gene Ontology terms, and spatial location. (B) Heatmap of top differentially expressed genes for each spatial domain. The color corresponds to the level of gene expression by z-score. (C) Top Gene Ontology Biological Process terms for each spatial layer ranked according to log-adjusted p-value. Dashed line represents the threshold for logP = 0.05. Bars are colored by total number of differentially expressed genes in each term. (D) Volcano plots of differentially expressed genes between young (left side) and middle-aged (right side) mice for each spatial domain. Genes were filtered by avg_log2FC < 0.5 and p_val_adj > 1e-3. Genes that passed both thresholds are colored according to their spatial domains.

Consistent with its location between the spared tissue and injury site epicenter, the scar interface region was characterized by expression of both white matter genes, such as *Plp1* and *Trf,* and immune cell genes such as *Cd36*, *Spp1*, and *Cd63* (Fig. 3B). Gene Ontology (GO) Biological Processes associated with the scar interface included wound healing as well as antigen processing and presentation (Fig. 3C). The necrotic and the infiltrating injury core regions were characterized by high expression of leukocyte-associated genes such as *Cd36*, *Thbs1*, *Pf4*, and *Ctsd* (Fig. 3B), but the necrotic region had significantly lower gene counts compared to other regions. The infiltrating injury core region was characterized by biological process associated with translation, myeloid leukocyte migration, and phagocytosis, whereas the necrotic region was associated with translation (Fig. 3C). The stromal layer showed the most distinct gene expression profile, being enriched for collagens, such as *Col1a1* and *Col8a1*, and other genes associated with fibrosis such as *Smoc2* (Fig. 3B). Accordingly, the stromal region was associated with the biological processes extracellular matrix organization, collagen fibril organization, and translation. This layer was present in the most superficial portion of the dorsal spinal cord, including regions that resemble the meninges, consistent with previous studies showing that fibrosis is not prevalent at 3 days post-SCI [26]. Spatial regions were then pseudobulked by region and age group, and each region was compared in gene expression between young and middle-aged mice. This comparison revealed very few genes (no more than six) that were differentially expressed between the two age groups (Fig. 3D).

### scRNA-seq reveals similar broad cell types between young and middle-aged mice acutely after SCI

To investigate the effect of middle-age on transcriptional changes and heterogeneity at the cellular level, we performed single cell RNA-seq (scRNA-seq) in young and middle-aged mouse injury sites at 3 dpi. These data were combined with our previous dataset [16] at the matching time points for a total of 4 young uninjured, 3 middle-aged uninjured, 3 young SCI, and 3 middle-aged SCI biological replicates in the combined dataset. There was a total of 71,794 cells, including 6,945 macrophages, 3,830 monocytes, 19,838 microglia, 12,940 endothelial cells, 1,304 tip cells, and 2,028 astrocytes (Supp. Fig. 6A). Dimensional reduction analysis of all the cells using UMAP showed distinct clusters that were uniquely identified using SingleR as well as manual annotation (Fig. 4A). We captured all major cell types as in our previous study except for oligodendrocytes. As we did not get middle-aged oligodendrocytes, they could not be compared with young oligodendrocytes and were excluded from further analyses. UMAP also showed that cells from young and middle-aged groups overlapped quite well, even without correcting for age or sample, indicating that the broad cell types are globally quite similar in gene expression profiles between the two age groups (Fig. 4B). As expected, there were clear separations in pre- and post-injury clusters (Fig. 4C). To confirm the identities of these broad cell clusters, we assessed the expression level of the top differentially expressed genes (DEGs) and annotated marker genes for each group (Fig. 4D). The 17 different cell types expressed the expected canonical marker genes. Many were unique to each cell type, but some were overlapping especially between myeloid cells that share similar lineage. The expression profile of most marker genes was similar between young and middle-aged groups with the exception of proliferation genes (e.g. *Mki67, Top2a*) in the Dividing Myeloid cluster that were higher in the middle-aged group. However, as noted above, this difference did not result in differences in myeloid proliferation between the two groups (Fig. 2A-C, Supp. Fig. 3A,C). After identification of each broad cell type using annotated marker genes, we separated them into myeloid, neural, and vascular categories for further subpopulation analysis.

**Figure 4.**
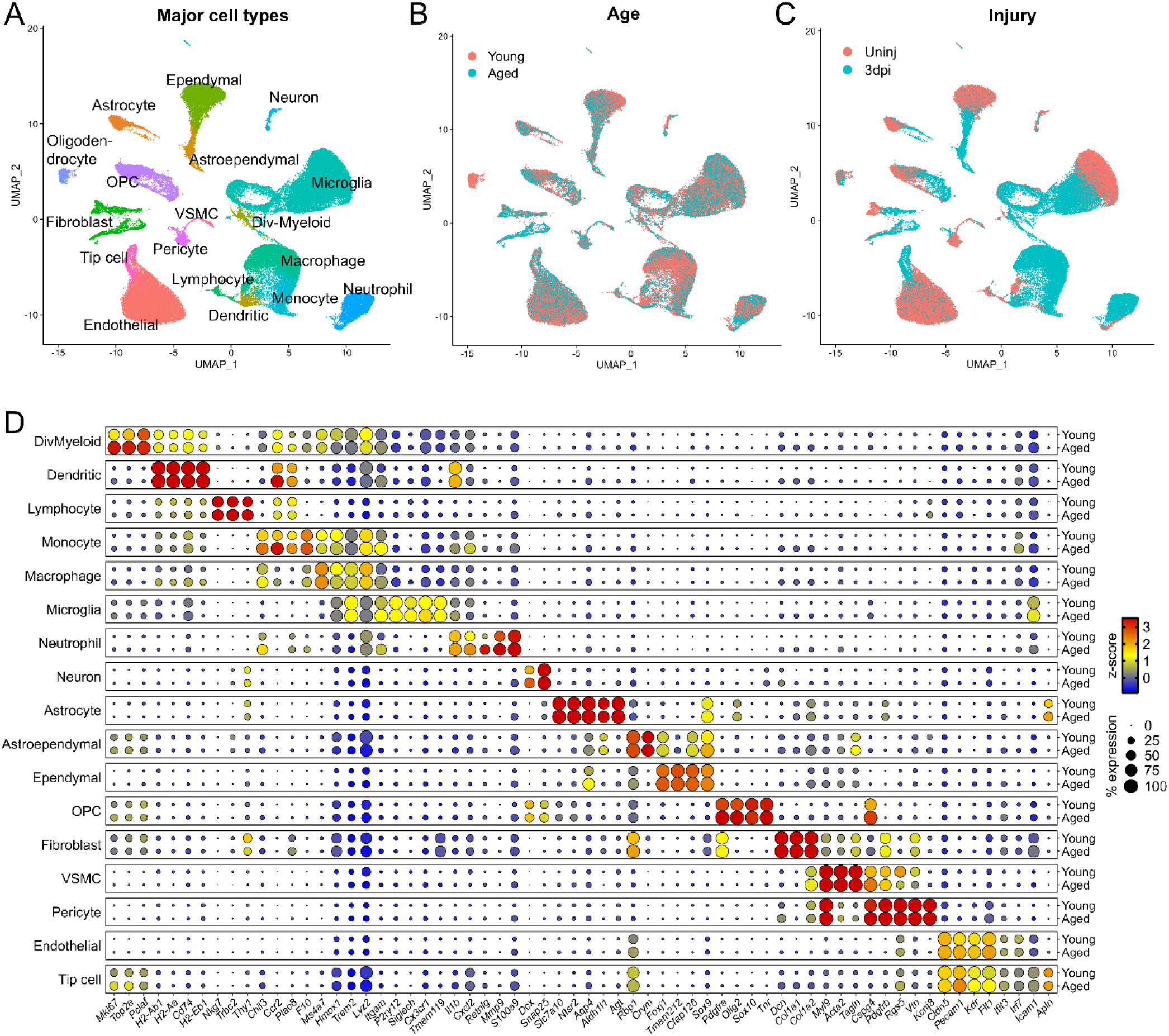
Single cell RNAseq analysis of the injury site from young and middle-aged mice show similar clustering and marker genes for general cell populations. (A) UMAP representing all cells from uninjured and 3 dpi spinal cords of young and aged mice, colored by cell type. A total of 4 young uninjured, 3 middle-aged uninjured, 3 young SCI, and 3 middle-aged SCI biological replicates represent the combined dataset for a total of 71,794 cells, including 6,945 macrophages, 3,830 monocytes, 19,838 microglia, 12,940 endothelial cells, 1,304 tip cells, and 2,028 astrocytes (see Supp. Fig. 6 for full list). Cells are annotated using canonical markers and alignment with other single cell references using SingleR. UMAP of all cells colored by age (B) or injury condition (C). Cells are shuffled in depth to show the distribution and density in localization in the UMAP. (D) Dotplot of marker genes used to annotate the general cell types. Circle size corresponds to the percentage of cells in the group which express at least one count of the gene. The color of the circle corresponds to the level of gene expression by z-score.

### scRNA-seq shows a different macrophage and neutrophil response in middle-aged mice

As one of the major pathobiologies in acute spinal cord injury, we first investigated subpopulation differences in immune cells and their transcriptional changes at the single-cell level. UMAP of the myeloid cells showed three large clusters pertaining to microglia, monocytes/macrophages, and neutrophils (Fig. 5A, Supp. Fig. 3A). Microglia were further clustered into homeostatic, and two types of SCI-induced microglia, Microglia-A (Micro-A) and Microglia-B (Micro-B). Monocytes gave rise to two macrophage populations, Macrophage-A (Macro-A) and Macrophage-B (Macro-B). This monocyte/macrophage cluster was also closely associated with dendritic cells, dividing myeloid cells, and border-associated macrophages (BAMs). The neutrophil cluster was comprised of homeostatic and activated subpopulations as noted by their presence in the uninjured and injured tissue respectively. Since an uninjured spinal cord does not normally contain neutrophils, this small population in our single cell data likely represents residual leukocytes in the vasculature and/or the leptomeninges after transcardial perfusion with artificial CSF. Without correction for age or batch effect the populations are visually well-integrated and evenly dispersed in lower-dimensional space indicating high homogeneity between the two age groups (Fig. 5B). As expected, there were injury-specific myeloid clusters (Fig. 5C). After injury, there is a reduction in the number of homeostatic microglia with a concomitant increase in Microglia-A and Microglia-B subpopulations in both young and middle-aged mice (Fig. 5D, Supp. Fig. 3B). Geno Ontology (GO) Biological Processes terms based on DEGs comparing young homeostatic microglia and young Microglia-A (Supp. Fig. 3C) were similar to those comparing the middle-aged counterparts (Fig. 5E). The top GO Biological Processes in Microglia-A from both age groups pertained to ATP synthesis, electron transport chain, and oxidative phosphorylation. Thus, the acute effects of SCI on microglia state seemed to be similar between young and middle-aged mice.

**Figure 5.**
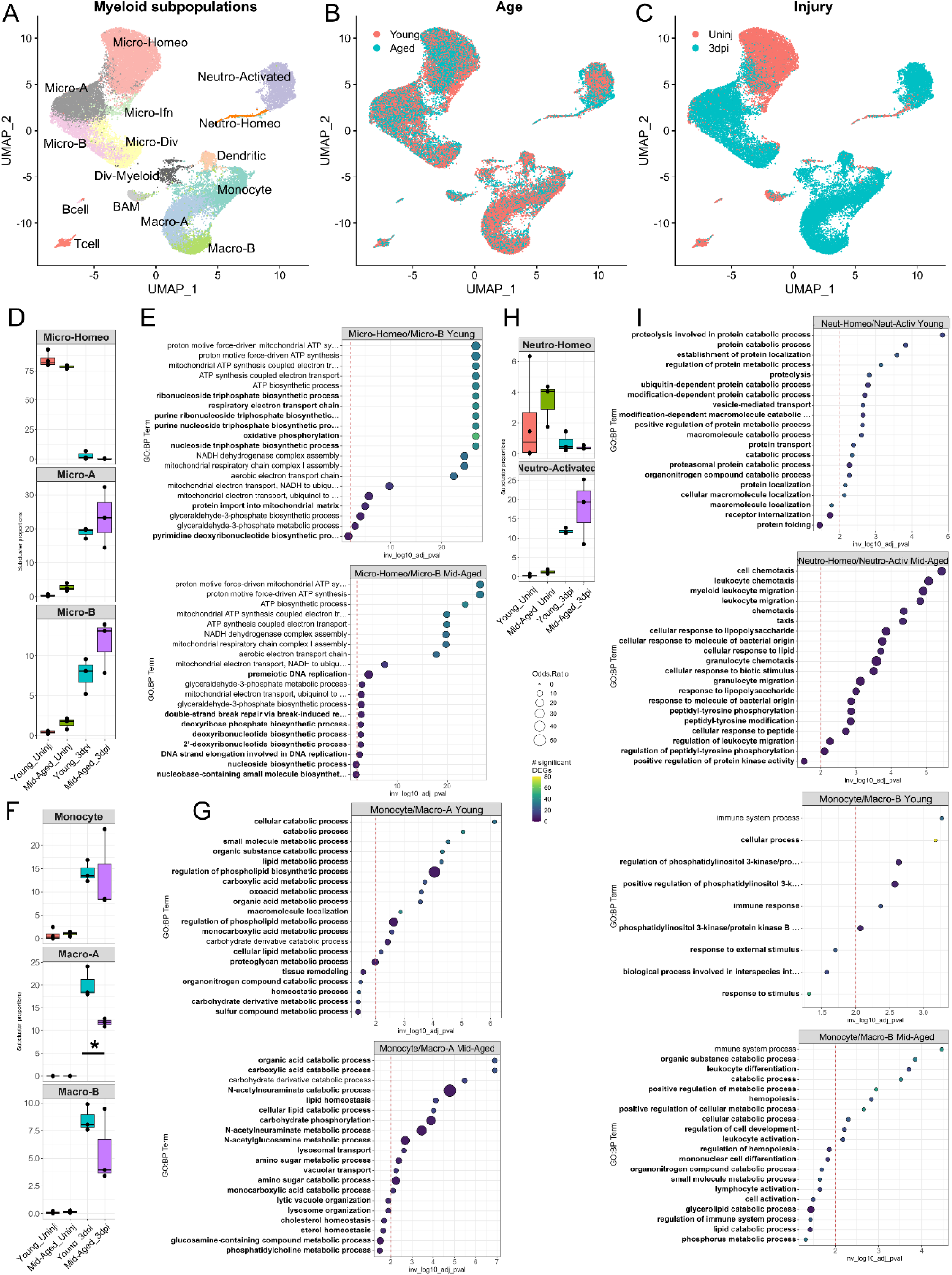
Single cell RNA-seq analysis of myeloid subpopulations in young and middle-aged mice after acute SCI. (A) UMAP of leukocyte subpopulations present in uninjured and 3 dpi spinal cords of young and middle-aged mice. Subpopulations are annotated using canonical markers and alignment with other single cell references using SingleR. (B) UMAP of leukocyte subpopulations separated by age (B) or injury condition (C). Cells are shuffled in depth to visualize the distribution and density in localization in the UMAP. Boxplots of microglia (D), monocyte/macrophages (F), neutrophil (H) subpopulation proportions that are shown as a percentage of all myeloid cells per-sample. Percentages are individually calculated for each sample and each spot represents one biological replicate. Boxplots are split by cell subtype with independent Y axes. Cell type proportions were statistically compared with ANOVA and Tukey’s post-test using a threshold of p<0.05. Gene Ontology Biological Processes terms based on differentially expressed genes between the noted subpopulations of microglia (E), monocytes/macrophages (G), and neutrophils (I) within their respective age groups. Micro-B was compared to Micro-Homeo, Macro-A and Macro-B were compared to monocytes, and Neutro-Activated was compared to Neutro-Homeo all within their respected age groups. Terms which are unique to each age group within a cellular response are bolded. Circle size represents the odds ratio and color represents the number of genes which map to each term. Volcano plots of DEG results are located in supplemental figure 3. Abbreviations -Micro-Homeo, Homeostatic Microglia; Micro-A, Microglia-A; Micro-B, Microglia-B; Micro-Ifn, Interferon-associated microglia; Micro-Div, Dividing Microglia; BAM, Border-associated Macrophages; Macro-A, Macrophage-A; Macro-B, Macrophage-B; Neutro-Activated, Activated Neutrophils; Neutro-Homeo, Homeostatic Neutrophils

Monocytes also gave rise to two macrophage populations, with Macrophage-A being the larger of the two (Fig. 5F). Interestingly, both Macrophage-A (p < 0.05) and Macrophage-B subpopulations were reduced in middle-aged mice, although the latter failed to reach statistical significance (Supp. Fig. 3B). GO Biological Processes based on DEGs between young monocytes and young Macrophage-A were largely like those comparing the middle-aged counterparts. The top GO Biological Processes terms in Macrophage-A from both age groups pertained to lipid and organic acid metabolism (Fig. 5G). However, young Macrophage-A were more associated with lipid metabolism while middle-aged Macrophage-A were more associated with organic acid metabolism. In addition, N-acetylneuraminate and N-acetylglucosamine metabolism were processes unique to middle-aged Macrophage-A subpopulation. GO terms for Macrophage-B were also different between young and middle-aged mice. While PI3K signaling was the major biological process in young Macrophage-B, it did not appear among the terms in middle-aged Macrophage-B, which was characterized by terms associated with catabolic and metabolic processes (Fig. 5G). Thus, the acute effects of SCI on monocyte-derived macrophages seem to be different between young and middle-aged mice.

Of the myeloid cells, neutrophils showed perhaps the starkest differences in GO Biological Processes between young and middle-aged mice. While comparison of young homeostatic neutrophils and young activated neutrophils resulted in GO Biological Processes pertaining to protein catabolism and transport, the corresponding comparison in middle-aged neutrophils resulted in terms associated with leukocyte migration and chemotaxis (Fig. 5I). This difference may explain the greater percentage of activated neutrophils observed in middle-aged mice, although it was not statistically significant (Fig. 5H). Taken together, scRNA-seq analysis of myeloid cells shows subpopulation-specific effects in macrophages and neutrophils, but not microglia, in middle-aged mice acutely after SCI.

### scRNA-seq shows an increased reactive OPC response in middle-aged mice

UMAP of neural cells showed three large clusters of ependymal cells, astrocytes, and oligodendrocyte progenitor cells (OPCs) (Fig. 6A). The few neurons that survived the dissociation procedure were classified as inhibitory, excitatory, and CSF-contacting neurons. Oligodendrocytes were mostly from our previous dataset [16] and were not present in the current experiment. Thus, neurons and mature oligodendrocytes were not analyzed further. UMAP showed three types of ependymal subclusters (A, B, and UK), but their marker gene expression showed few differences (Supp. Fig. 4A), so they were combined into one group in downstream analysis. These ependymal cells were the predominant subcluster in the uninjured spinal cord, whereas SCI gave rise to astroependymal-A and astroependymal-B subclusters, whose proportions were not significantly different between young and middle-aged mice (Fig. 6D, Supp. Fig. 4B). GO Biological Processes terms based on DEGs comparing young ependymal cells and young astroependymal-A cells were similar to those comparing the middle-aged counterparts (Fig. 6E). The top GO Biological Processes in astroependymal-A from both age groups pertained to complement activation and humoral immune response activation. GO Biological Process in astroependymal-B cells were also similar in young and middle-aged mice, except for young mice showing enrichment for genes related to mitochondrial function (Fig. 6F).

**Figure 6.**
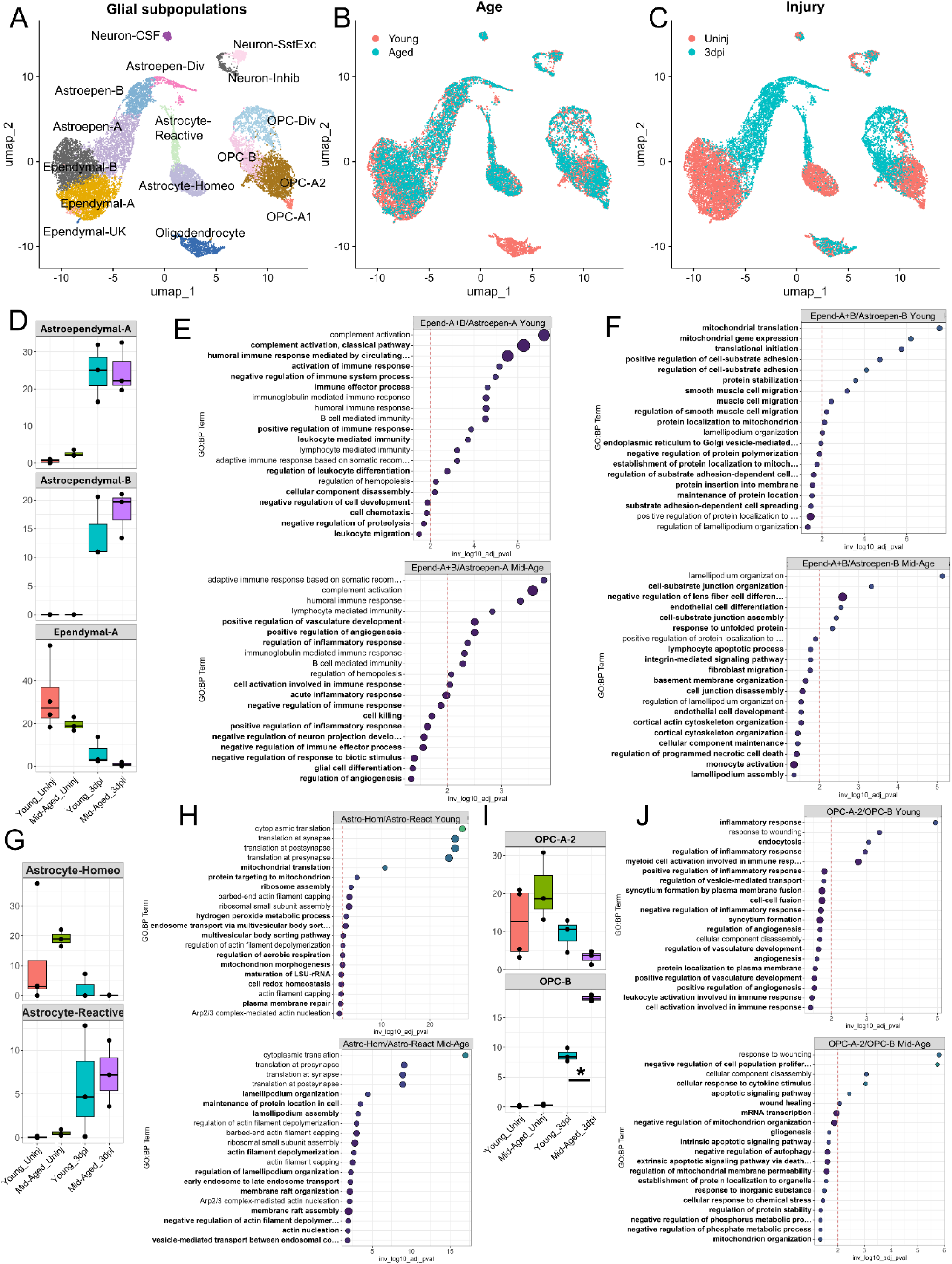
Single cell RNA-seq analysis of glial subpopulations in young and middle-aged mice after acute SCI. (A) UMAP of glial subpopulations present in uninjured and 3 dpi spinal cords of young and middle-aged mice. Subpopulations are annotated using canonical markers and alignment with other single cell references using SingleR. (B) UMAP of glial subpopulations separated by age (B) or injury condition (C). Cells are shuffled in depth to visualize the distribution and density in localization in the UMAP. Boxplots of ependymal cell subpopulations (D) and their Gene Ontology Biological Processes based on DEG comparing combined ependymal cells A and B to the astroependymal A subpopulation (E) or the astroependymal B subpopulation (F). Boxplots of astrocyte subpopulations (G) and their Gene Ontology Biological Processes based on DEG comparing homeostatic and reactive astrocytes (H). Boxplots of oligodendrocyte progenitor cell (OPC) subpopulations (I) and their Gene Ontology Biological Processes based on DEG comparing OPC-a2 and OPC-b (J). Boxplot percentages are individually calculated for each sample and each spot represents one biological replicate. Boxplots are split by cell subtype with independent Y axes. Cell type proportions were statistically compared with ANOVA and Tukey’s post-test using a threshold of p<0.05. GO terms are from comparisons of cell types within their respective age group. Terms which are unique to each age group within a cellular response are bolded. Circle size represents the odds ratio and color represents the number of genes which map to each term. Volcano plots of DEG results are located in supplemental figure 4. Abbreviations – Astroepen. Astroependymal; Astroepen-Div, Astroependymal-Dividing; Astrocyte-Homeo, Astrocyte-Homeostatic; Neuron-CSF, Cerebrospinalfluid-contacting Neurons; Neuron-SstExc, Sst positive and Excitatory Neurons; Neuron-Inhib, Inhibitory Neurons; OPC, Oligodendrocyte Precursor Cell; OPC-Div, Oligodendrocyte Precursor Cell-Dividing

The astrocyte cluster was comprised of a subcluster predominant in uninjured spinal cord (i.e. homeostatic) and a subcluster predominant in the injured spinal cord (i.e. reactive), which likely represents the progression of astrocytes during astrogliosis (Fig. 6C). Although 3 dpi is toward the early phase of astrogliosis, the scRNA-seq analysis was able to identify a reactive subpopulation characterized by increased expression of *Gfap* and *Osmr* (Supp. Fig. 4A). However, the proportion of the two subpopulations as well the biological processes defining their transition from homeostatic to reactive astrocytes were similar between young and middle-aged mice (Fig. 6G-H). Most of the top biological processes that pertained to reactive astrogliosis at this early time point were related to translation and ribosomal assembly.

The OPC cluster was comprised of a homeostatic subpopulation (OPC-A) predominant in the uninjured spinal cord, and two subpopulations that appeared after SCI, one of which we classified as dividing OPCs (OPC-div) due to their unique expression of cell cycle genes (Fig. 6A-C, Supp. Fig. 4A). The other injury-specific OPC subpopulation (OPC-B) that likely represents a reactive phenotype was significantly higher (p < 0.05) in the middle-aged group seemingly at the expense of the homeostatic OPC-A subpopulation (Fig. 6I). In addition, biological processes comparing OPC-A and OPC-B subpopulations from the respective age groups showed interesting differences. Whereas young reactive OPCs (OPC-B) were characterized by inflammatory responses compared to young homeostatic OPCs (OPC-A), these inflammatory processes were largely absent in the middle-aged OPCs (Fig. 6J). One biological process that was common to both ages groups was response to wounding. Taken together, at 3 days post-SCI, astrocyte and OPC lineage cells seem to display similar response to SCI in both young and middle-aged mice, but reactive OPCs seem to be greater in proportion as well as less inflammatory in middle-aged mice.

### Vascular cell activation after SCI is broadly comparable between young and aged mice

UMAP of vascular cells displayed four large clusters comprised of an endothelial cluster, pericyte/vascular smooth muscle cell (VSMC) cluster, and two distinct fibroblasts clusters (Fig. 7A-C). The endothelial cluster was comprised of arterial, venous, three capillary, and one tip cell subpopulations. All subpopulations expressed classic endothelial markers such as *Pecam1* and *Cldn5* (Supp. Fig. 5A). The three capillary groups appear to broadly represent a gradient in transition from arterial to venous populations, however, these three groups did not strongly express unique genes or ontologies and so were combined for downstream analyses. Arterial cells were greatly reduced after injury in both middle-aged and young populations, comprising a primarily homeostatic population, whereas venous cells increased after injury (Fig. 7D), expressing elevated levels of multiple cell adhesion molecules (*Vcam1*, *Selp,* etc.). The GO Biological Processes terms for the injured arterial cells showed immune response in young mice, but cell adhesion and vascular development in middle-aged mice (Fig. 7E). Tip cells were almost exclusively dominated by proliferation-associated genes in both groups, with younger tip cells having mitochondrial translation and mitochondrial gene expression as the top two terms (Fig. 7E).

**Figure 7.**
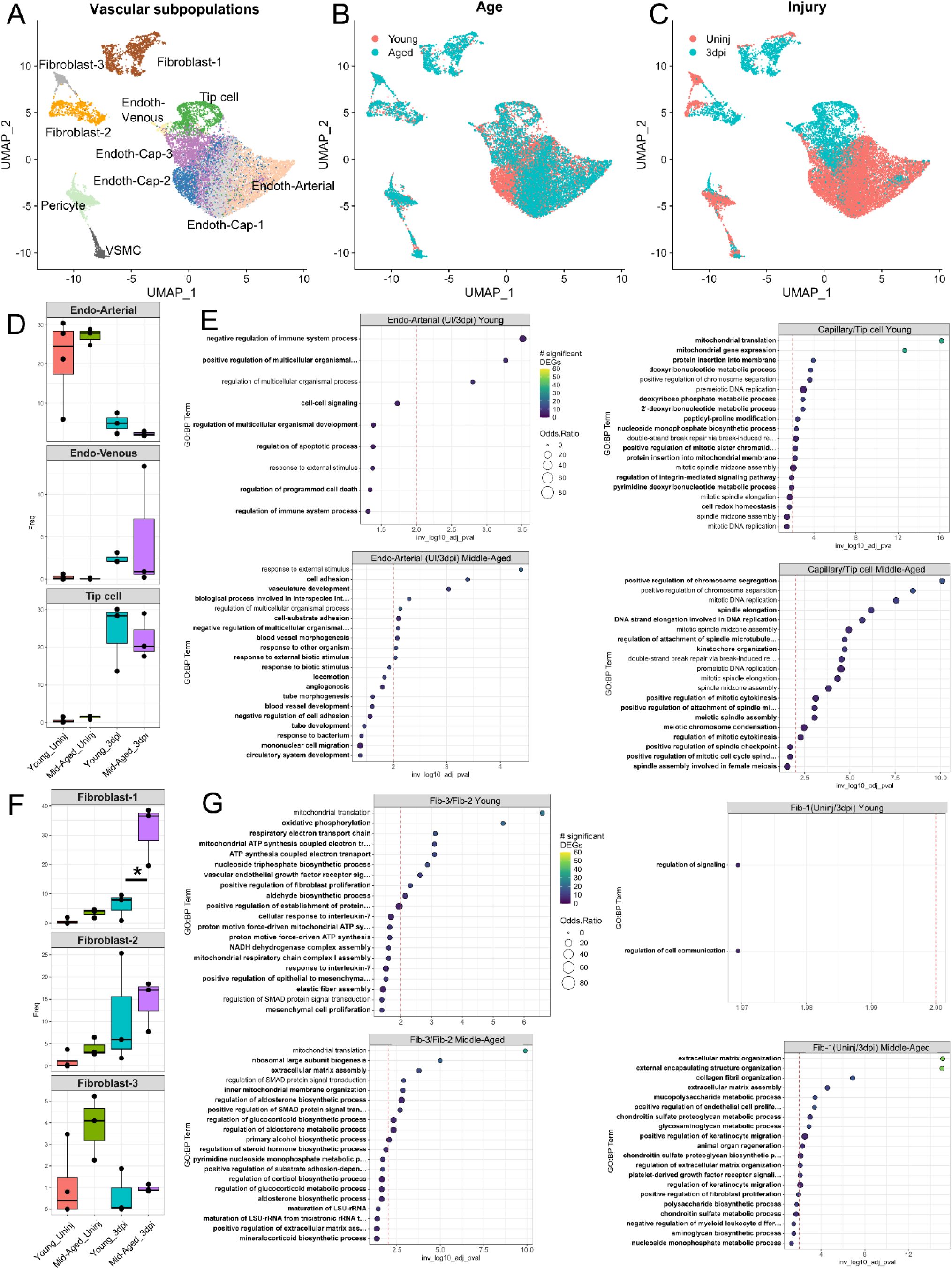
Single cell RNA-seq analysis of vascular subpopulations in young and middle-aged mice after acute SCI. (A) UMAP of vascular subpopulations present in uninjured and 3 dpi spinal cords of young and middle-aged mice. Subpopulations are annotated using canonical markers and alignment with other single cell references using SingleR. UMAP of vascular subpopulations separated by age (B) or injury condition (C). Cells are shuffled in depth to visualize the distribution and density in localization in the UMAP. Boxplots of endothelial subpopulations (D) and their Gene Ontology Biological Processes based on DEG comparing arterial uninjured and injured or capillary to tip cells (E). Boxplots of fibroblast subpopulations (F) and their Gene Ontology Biological Processes based on DEG comparing fibroblast-3 to fibroblast-2 or uninjured fibroblast-1 to injured fibroblast-1 (G). Boxplot percentages are individually calculated for each sample and each spot represents one biological replicate. Boxplots are split by cell subtype with independent Y axes. Cell type proportions were statistically compared with ANOVA and Tukey’s post-test using a threshold of p<0.05. GO terms are from comparisons of cell types within their respective age group. Terms which are unique to each age group within a cellular response are bolded. Circle size represents the odds ratio and color represents the number of genes which map to each term. Volcano plots of DEG results are located in supplemental figures 5. Abbreviations – Endoth, Endothelial; Cap, Capillary; VSMC, Vascular Smooth Muscle Cell

The fibroblast populations consisted of three groups split between two clusters in the UMAP, with group 2 and 3 forming one cluster and group 1 forming another. Group 3 consists of mostly uninjured fibroblasts and expresses the highest levels of genes associated with homeostatic perivascular fibroblasts (*Dcn*, *Lum*) while group 2 consists mostly of injured fibroblasts with reduced homeostatic markers (Supp. Fig. 5A). Both groups show an increase in populations in response to injury, with group 2 contributing to about 15 percent of the vascular population in the aged group, and 5 percent in the young group (Fig. 7F; Supp. Fig. 5B). Fibroblast group 1 expresses the highest level of collagens and *Postn*, associated with fibroblast activation and ECM deposition (Supp. Fig. 5A, C). Interestingly, group 1 also is enriched in various solute channels, a marker of human meningeal fibroblasts in a recent study [27]. Group 1 also showed the most dramatic increase with a significant difference between age groups (Supp. Fig. 5B). For fibroblast group 1, only the middle-aged group showed significant GO biological terms with their differentially expressed genes when comparing uninjured to 3 dpi cells (Fig. 7G). These terms were primarily related to ECM organization. Fibroblast-2 was compared to Fibroblast-3 because they clustered together on the UMAP with the former present in the injured tissue, and the latter present in the uninjured tissue, suggesting that Fibroblast-2 was arising from Fibroblast-3 after SCI (Fig. 7B-C; Supp. Fig. 5B). Fibroblast-2 GO terms in the young were associated with oxidative phosphorylation and ATP synthesis, whereas GO terms in the middle-aged were associated with mitochondrial membrane, ECM assembly, and SMAD signaling (Fig. 7G). Taken together, although we did not find vascular subpopulations unique to young or middle-aged mice, the transcriptional responses for many subpopulations were different between the two age groups.

## Discussion

The goal of this study was to use spatial and single cell transcriptomics to investigate cellular and molecular differences between young (2-4 months) and middle-aged (10-12 months) mice after contusive SCI. In contrast to previous studies, histological analysis revealed similar lesion areas between the two age cohorts, with no significant differences in the density of lesional myeloid cells, fibroblasts, or astrocytes, and no marked differences in cell proliferation between them. Behaviorally, both groups exhibited comparable locomotor recovery, with no significant differences in Basso Mouse Scale (BMS) scores or subscores following SCI. Our spatial transcriptomic analysis corroborated the gross histological similarity between molecular responses in both groups, further indicating that middle-aged mice do not show significantly impaired pathology compared to younger counterparts following SCI. Our single cell transcriptional data, however, showed that subpopulations of myeloid, glial, and vascular cells have altered proportions and predicted alterations in function with age. Taken together, these findings suggest that progression from youth to middle-age may not significantly affect the immediate cellular and functional recovery from SCI, but that effects of age may be manifested at the transcriptional level during middle-age before becoming pathological later in life.

Despite observing single-cell differences in immune and fibrotic responses in middle-aged animals, we did not detect any corresponding behavioral or histological deficits—contrary to our initial hypothesis. While previous studies have reported age-related impairments in locomotor recovery following similar injuries and postoperative care, we found no significant differences in open-field locomotor recovery between our young (2–3 months) and middle-aged (10–12 months) mice. One possible explanation is that our middle-aged mice were slightly younger (by 1–2 months) than those used in earlier studies. Prior research has typically examined mice aged 14 to 18 months [8, 9, 15, 28] which roughly corresponds to humans just over 40 years old—a stage linked to a period of rapid biological aging [4, 29].

Prior studies examining old mice (18-24 months) in spinal cord injury models have reported immune alterations that are associated with behavioral deficits [15]. In our study, while middle-aged mice exhibited an increased number of activated microglia during the acute phase, this alone does not seem to be sufficient to cause functional impairments and further immune dysregulation, as observed in old mice, may be required to produce behavioral deficits. Interestingly, we also observed a reduction in macrophage number in middle-aged mice, consistent with findings by Salvador et al., who reported a more pronounced disparity between young and old mice [15]. It is intriguing to speculate that this relative reduction in early monocyte recruitment and differentiation may contribute to long-term deficits in locomotor recovery. Conversely, these findings suggest that enhancing monocyte recruitment or accelerating their differentiation in aged animals could potentially improve recovery outcomes.

Another major difference between young and middle-aged mice was the presence of PI3K signaling as a top GO term in young, but not middle-aged, macrophage-b subpopulation. Interestingly, we recently reported that PI3K activation leads to accumulation lipids in foamy macrophages that appear at the spinal cord injury site, which contributes to their pro-inflammatory state [30]. The absence of PI3K GO terms in middle-aged mice would suggest that there should be less lipid accumulation in middle-aged macrophages, which would be predicted to have beneficial effects. However, it could also mean reduced phagocytic abilities in middle-aged macrophages that could be associated with insufficient debris clearance. This difference in PI3K signaling between young and middled aged mice requires further investigation in the future.

OPCs have recently been characterized as contributing directly to inflammation by antigen presentation and expression of cytokines [31–33]. Consistent with these studies, the top GO terms for OPC-b in young mice after SCI were related to inflammation, but surprisingly, these inflammation GO terms disappeared in middle-aged OPC-b cells. The functional consequence of OPC-related inflammation seems to be context-dependent with both detrimental and beneficial roles. Activated OPCs can damage the blood-brain barrier and present antigen to promote inflammation [33, 34]. On the other hand, OPCs can regulate inflammation to activate the neuroprotective effects of astrocytes and microglia [35]. In the context of our studies, we speculate that the absence of the beneficial OPC-associated inflammation may contribute to the worse outcomes in old mice, but further studies are needed to test this hypothesis.

Although there were no histological differences in the fibrotic scar between young and middle-aged mice, our single cell data showed increased presence of the fibroblast-1 subpopulation in middle-aged mice after SCI. Recent studies have shown that fibroblasts represent several distinct subsets in homeostatic and injury states with distinct spatial and transcriptional profiles [26, 27, 36–38]. Interestingly, fibroblast-1 expressed markers of meningeal fibroblasts and GO terms associated with this subpopulation in middle-aged mice were related to ECM deposition and organization. This raises the possibility that fibrosis in middle-aged mice may lead to greater ECM deposition. As these fibroblasts are localized with macrophages at the injury site, changes in the ECM environment could lead to altered macrophage polarization states as previously described [39, 40]. Alternatively, the altered macrophage types and/or states as described in our data above could be differentially recruiting the fibroblast subpopulations [17].

Sex as a biological variable has been shown to have significant impacts on outcomes in spinal cord injury for both humans and mice [41] and the majority of SCI patients are male (NSCISC database). One limitation of this study is the exclusive use of female mice. Female mice were chosen due to the low number of biological replicates able to be processed for spatial transcriptomics, and for the ability to combine the new sequencing dataset with our previous one. Future studies would greatly benefit from the use of males to determine if different transcriptomic changes are observed.

Another objective of this study was to provide a spatial and single-cell transcriptome resource for other researchers. With the addition of ∼42k cells over 8 samples in our new dataset focusing on 3 dpi, we were able to resolve multiple new cell subtypes associated with homeostatic and injured tissue that we were not able to observe in our previous study. For example, we were able to resolve uninjured and injury-associated neutrophil clusters, and distinct T-cell and B-cell lymphocyte clusters. In neural cells, we resolved homeostatic and reactive astrocytes, multiple astroependymal subtypes including a proliferating group, and captured enough neurons to form three broad clusters (of note is the CSF-contacting neurons). Lastly, in vascular cells, we resolved three distinct subpopulations of fibroblasts that showed injury- and age-dependent effects. A limitation to our dataset, however, is that batch effects precluded the analysis of oligodendrocytes as they were not obtained in the samples generated for this study. We also note that capillary group 2 appears to be primarily from our original dataset as well. As far as we are aware, this is the first dataset published to encompass nearly every cell type present in the acute phase of spinal cord injury for middle aged mice.

## Conclusions

In conclusion, our findings indicate that compared to young mice (2-4 months), SCI in middle-aged mice (10-12 months) show transcriptional differences in immune, glial, and fibrotic response at the single cell level. However, these differences did not manifest in locomotor recovery or histopathology as assessed by immunohistochemistry or spatial transcriptomics. Our results contrast with previous studies that used slightly older middle-aged mice (14-18 months), suggesting that middle-age is a period of rapid decline in health that may contribute to the large heterogeneity in pathology and recovery observed in people who sustain SCI. As the middle-age now represents the median age of incidence for SCI, it will become important to better understand the possible rapid health changes that occur in this population, and how this information can be used (e.g. through biomarkers) to reduce heterogeneity in SCI clinical trials.

## List of Abbreviations

BMS: Basso Mouse Scale
SCI: Spinal Cord Injury
GFAP: Glial Fibrillary Acidic Protein
CD11b: Cluster of Differentiation 11b
PDGFRβ: Platelet-Derived Growth Factor Receptor Beta
EdU: 5-ethynyl-2’-deoxyuridine
ECM: Extracellular Matrix
OPC: Oligodendrocyte Precursor Cell
GO: Gene Ontology
VSMC: Vascular Smooth Muscle Cell

## Declarations

### Ethics approval and consent to participate

All animal procedures were conducted in accordance with the guidelines established by the Institutional Animal Care and Use Committee (IACUC). Ethical approval was obtained for all animal experiments.

### Consent for publication

Not applicable

### Availability of data and materials

The datasets generated during and/or analyzed during the current study are available from the corresponding author on reasonable request. Sequencing data used in this study from previously published work can be found under GSE162610. Sequencing data generated by this study have been uploaded onto GEO and can be accessed upon request to the corresponding author while the manuscript is under review. It will be released to the public once the manuscript is accepted and become publicly available. Detailed animal and quantification data has been deposited in the ODC-SCI database.

### Competing interests

The authors declare that they have no competing interests.

### Funding

This work was supported by NINDS RM1NS133003, R01NS081040, R21NS123492, Else Kroner-Fresenius Foundation (2019-A54), The Buoniconti Fund to Cure Paralysis, The Miami Project to Cure Paralysis. DJ was supported by the John T. and Winifred Hayward Foundation Genomic Medicine Research Fellowship. JKL is the Christine E. Lynn Distinguished Professor in Neuroscience.

### Authors’ contributions

CF, DJ, BK, JSC, JKL designed and performed the experiments, collected and analyzed the results. SC and AB performed the experiments. CF, DJ, AB, JKL wrote the manuscript.

## Acknowledgements

We would like to thank Yan Shi at the Miami Project Imaging Core, and Maria Boulina at the Analytical Imaging Core Facility (S10OD023579) at the University of Miami School of Medicine. We would like to thank Bill Hulmes at the Hussman Institute for Human Genomics Sequencing Core. We would like to thank Shaffiat Karmally and Tatyana Camejo for their assistance with collecting BMS data.

---

**Supplemental Figure 1.**
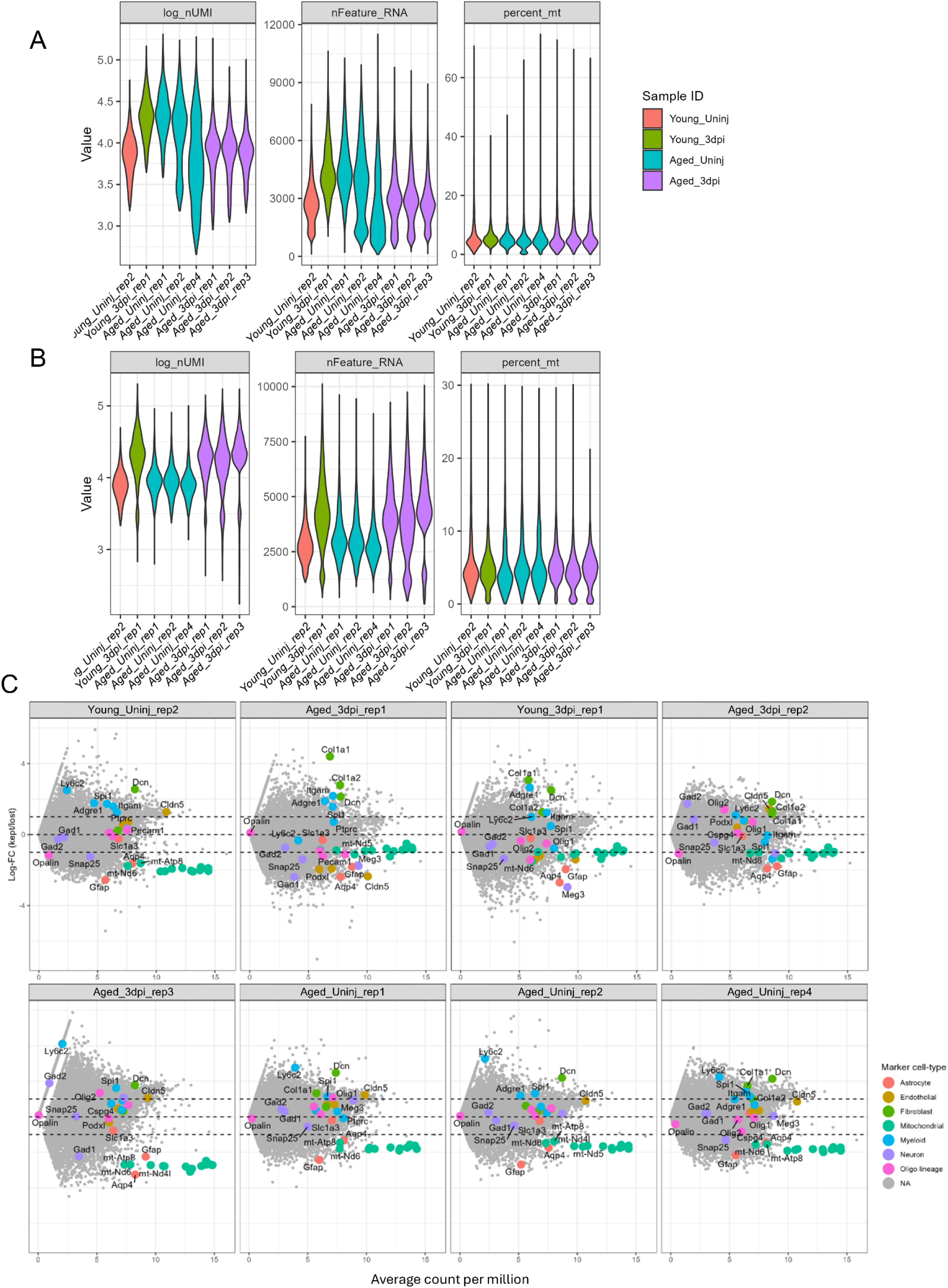
Quality control metrics for spots before filtering. Log numbers of UMIs, features, and percent mitochondrial counts per-droplet per-sample before (A) and after (B) quality control. (C) Comparison of gene expression between passing and removed droplets. Retained droplet genes represented as positive log fold-change, and removed drops are negative. Genes used to manually annotate cell types are colored. Dashed lines represent 0 and a p-value of 0.05. Note mitochondrial genes are enriched in the removed droplets in all samples.

**Supplemental Figure 2.**
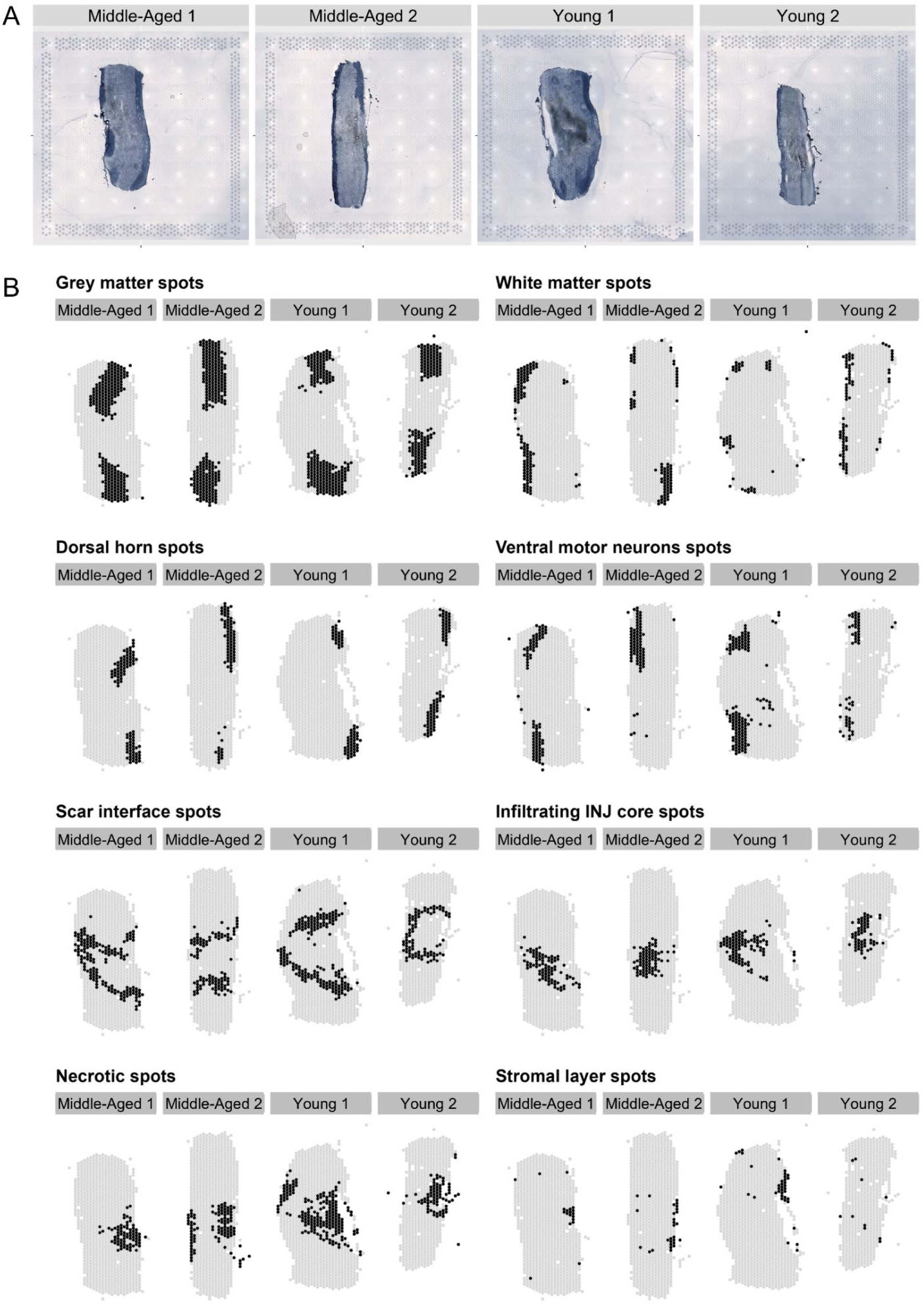
Visium spatial transcriptomics clusters in each biological replicate. (A) Sudan black stain of a 10X Visium slide with four different biological replicates. (B) Locations of the identified spatial clusters in each biological replicate.

**Supplemental Figure 3.**
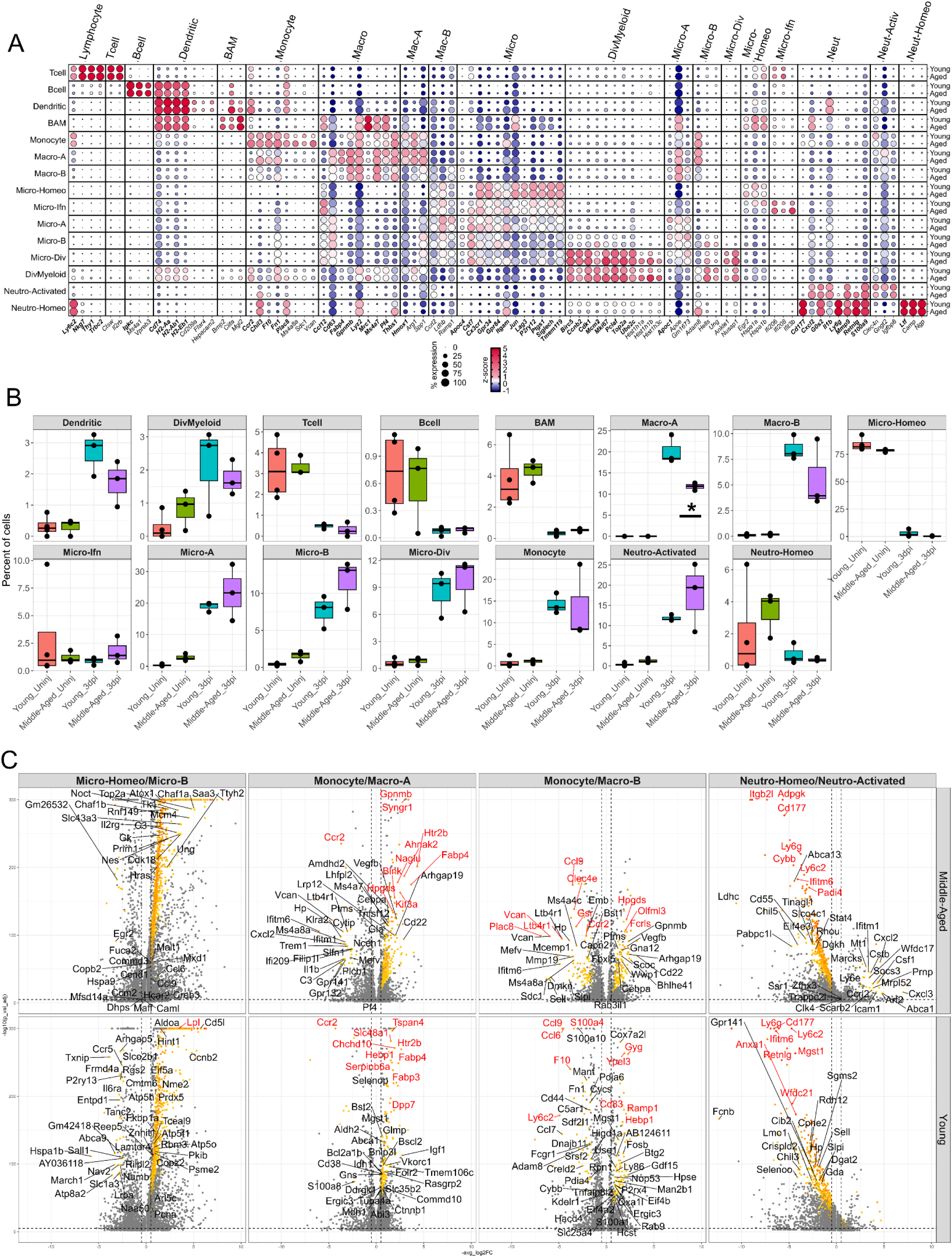
Leukocyte subpopulation marker gene expression and proportions. (A) Dotplot of genes used to manually annotate the leukocyte cell types (bold) and top genes found by differential gene expression analysis (not bold). Genes are binned by groups. Cell types are split by age. Circle size corresponds to the percentage of cells in the group which express at least one count of the gene. The color of the circle corresponds to the level of gene expression by z-score. (B) Boxplots of cell subpopulation proportions. Percentages are individually calculated for each sample and each data point represents one biological replicate. Cell type proportions were statistically compared with One-Way ANOVA and Tukey’s post-test. *p<0.05 between ages for each injury timepoint. (C) Volcano plots of differentially expressed genes from myeloid subpopulations. Genes with negative log2FC represent the homeostatic state and positive log2FC are the genes that are increased after injury. Genes are filtered to an adjusted p-value of less than or equal to 1e-5, an absolute value of average log2FC greater than or equal to 0.5, and a percent difference greater than 0.25. Genes which pass filtering are colored by percent difference. The top 10 genes weighted by percent difference (red) and genes that are unique between young and aged for each group (black) are annotated. Dashed lines represent log_2_FC and adjusted p-value thresholds

**Supplemental Figure 4.**
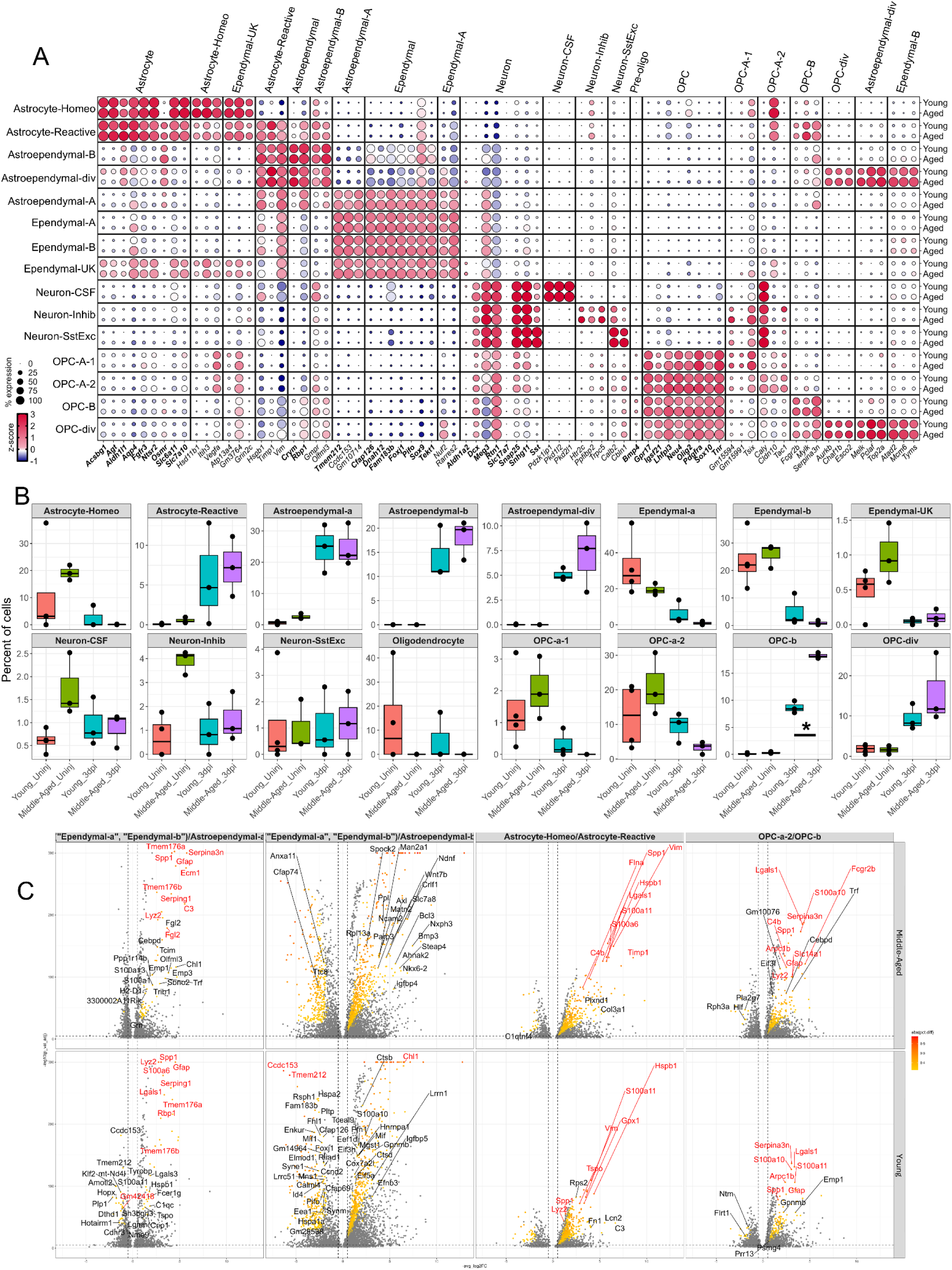
Glial cell subpopulation marker gene expression and proportions. (A) Dotplot of genes used to manually annotate the glial cell types in the study (bold) and top genes found by differential gene expression (not bold). Genes are binned by groups. Cell types are split by age. Circle size corresponds to the percentage of cells in the group which express at least one count of the gene. The color of the circle corresponds to the level of gene expression by z-score. (B) Boxplots of cell subpopulation proportions. Percentages are individually calculated for each sample and each spot represents one biological replicate. Boxplots are split by cell subtype with independent Y axes. Boxplots are colored by age and days post injury. Cell type proportions were statistically compared with One-way ANOVA and Tukey’s post-test. *p<0.05 between ages for each injury timepoint. (C) Volcano plots of gene expression changes after injury. Genes with negative log2FC represent the homeostatic state while positive log2FC represents those expressed in the injured state. Genes are filtered to an adjusted p-value of less than or equal to 1e-5, an absolute value of average log2FC greater than or equal to 0.5, and a percent difference greater than 0.25. Genes which pass filtering are colored by percent difference. The top 10 genes weighted by percent difference (red) and genes that are unique between young and aged for each group (black) are annotated. Dashed lines represent log2FC and adjusted p-value thresholds.

**Supplemental Figure 5.**
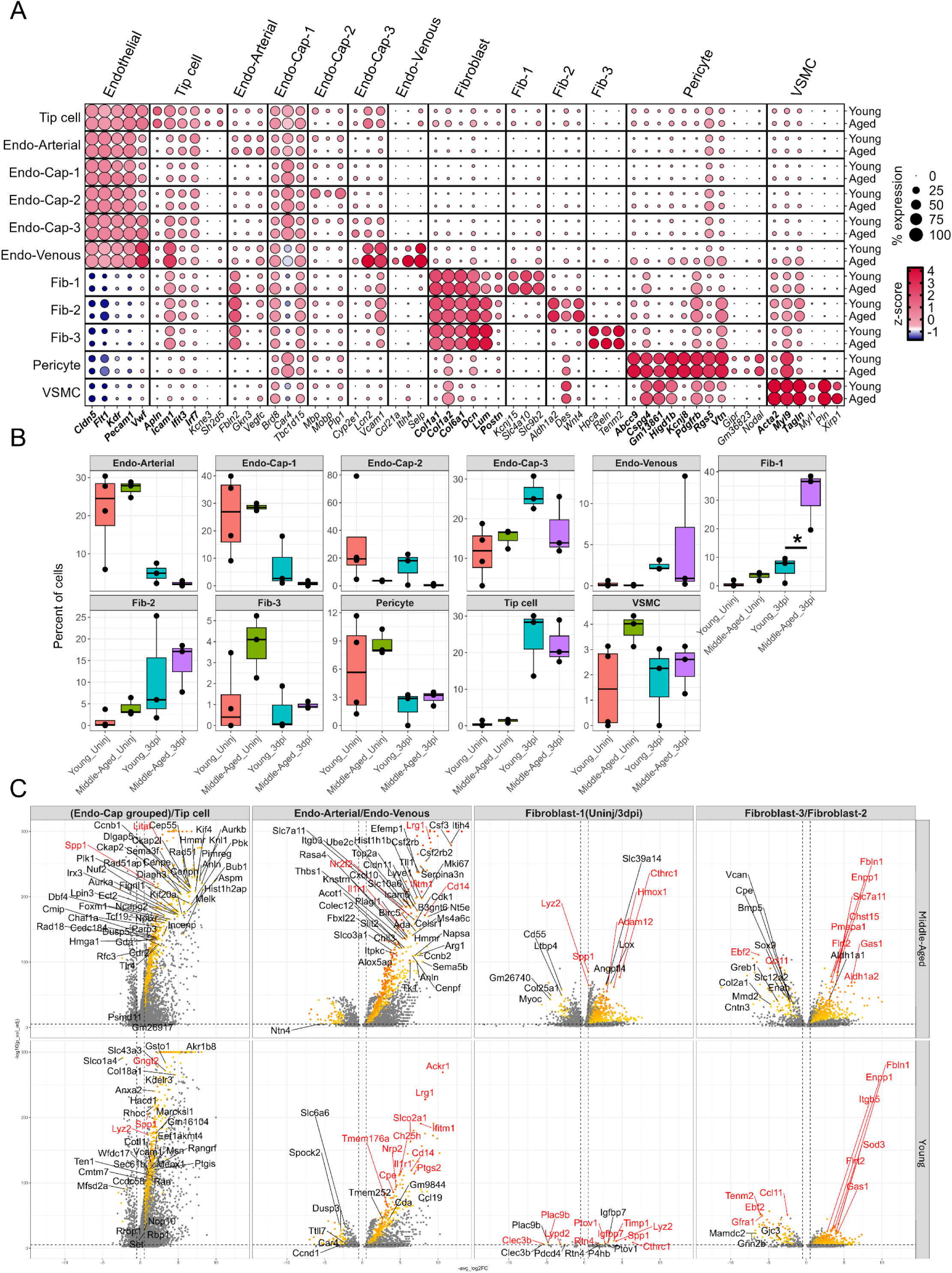
Vascular cell subpopulation marker gene expression and proportions. (A) Dotplot of genes used to manually annotate the vascular cell types in the study (bold) and top genes found by differential gene expression (not bold). Genes are binned by groups. Cell types are split by age. Circle size corresponds to the percentage of cells in the group which express at least one count of the gene. The color of the circle corresponds to the level of gene expression by z-score. (B) Boxplots of cell subpopulation proportions of each sample. Percentages are individually calculated for each sample and each spot represents one biological replicate. Boxplots are split by cell subpopulation with independent Y axes. Boxplots are colored by age and days post injury. Cell type proportions were statistically compared with One-way ANOVA and Tukey’s post-test. *p<0.05 between ages for each injury timepoint. (C) Volcano plots of transcriptional responses from different cell types to injury. Genes with negative log2FC represent the homeostatic state while positive are the genes increased after injury. Genes are filtered to an adjusted p-value of less than or equal to 1e-5, an absolute value of average log2FC greater than or equal to 0.5, and a percent difference greater than 0.25. Genes which pass filtering are colored by percent difference. The top 10 genes weighted by percent difference (red) and genes that are unique between young and aged for each group (black) are annotated. Dashed lines represent log2FC and adjusted p-value thresholds. Vascular cells without uniquely annotated pre- and post-injury cell subtypes were split and compared by uninjured and 3dpi populations.

**Supplemental Figure 6.**
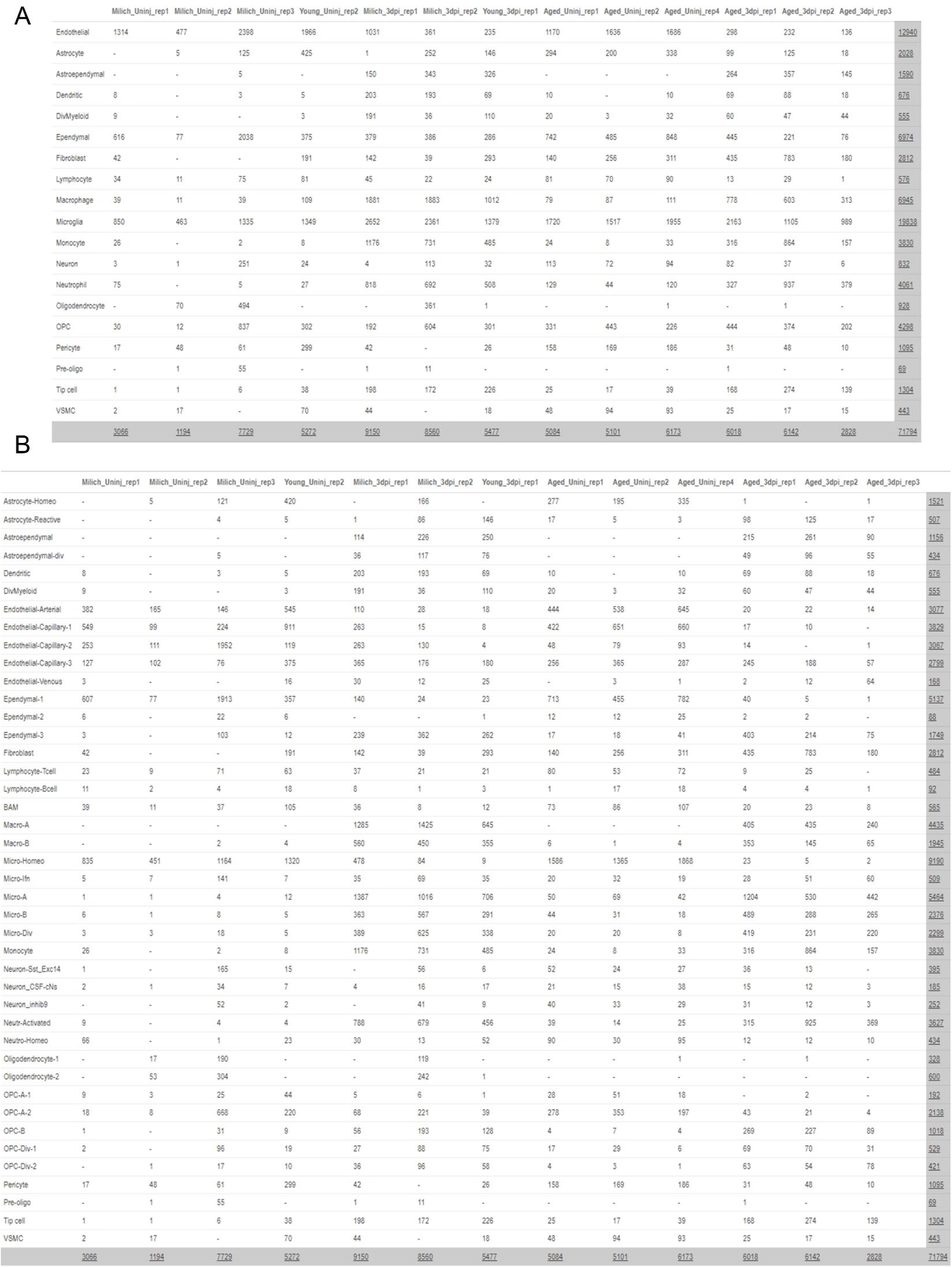
Table of cell counts obtained in each sample. (A) Table of cell counts per-sample for all broad cell types. (B) Table of cell counts per-sample for all cell subtypes. Row and column sums are highlighted.

## Notes

https://www.ncbi.nlm.nih.gov/geo/query/acc.cgi?acc=GSE162610

### Competing Interest Statement

The authors have declared no competing interest.

